# Carbohydrate adaptation drives liver-brain axis maturation

**DOI:** 10.1101/2025.11.05.685548

**Authors:** Hongmei Cui, Zheng Wu, Yuannyu Zhang, Hieu S. Vu, Hongli Chen, Xiaofei Gao, Yan Jin, Donghong Cai, Sarada Achyutuni, Phong Nguyen, Chunxiao Pan, Hui Cao, Camenzind G. Robinson, Jeffrey D. Steinberg, Laura J. Janke, Sara M. Nowinski, Jian Xu, Ralph J. DeBerardinis, Min Ni

## Abstract

Mammalian postnatal life requires adaptation to a carbohydrate-rich diet, yet how metabolic programs are coordinated within and across organs is unclear. Using time-resolved transcriptomic and metabolomic analyses from the neonatal period through adulthood, we show that mouse liver rapidly acquires oxidative and detoxification capacity after weaning. This transition enables the brain to establish energy-sufficient, low-toxicity metabolic environment for neuronal development. This maturation process is marked by progressive activation of the hepatic electron transport chain (ETC), with the mitochondrial RNA endoribonuclease LACTB2 acting as a key regulator. LACTB2 prevents the accumulation of mitochondrial RNAs and sustains expression of mtDNA-encoded ETC subunits, thereby preserving mitochondrial competence for oxidative metabolism. LACTB2 is postnatally induced in hepatocytes, and its loss causes defective glucose utilization, systemic metabolic toxicity, and impaired brain metabolism and myelination, leading to prepubertal lethality, particularly in males. Restoring ETC function through liver-targeted expression of yeast NADH dehydrogenase NDI1, inhibiting the integrated stress response or ammonia scavenging improved survival. Our findings identify LACTB2-dependent hepatic mitochondrial maturation as a central mechanism that aligns carbohydrate adaptation with the liver-brain metabolic coordination to support early-life development.

## INTRODUCTION

Mammalian development requires metabolic flexibility to support rapid growth and organ maturation^1,2^. A key challenge in early life is the nutritional shift from fat- and protein-rich milk to carbohydrate-based foods after weaning^3^. This transition necessitates the establishment of glucose oxidation, lipid metabolism, nitrogen disposal, and detoxification pathways, requiring the liver’s maturation from a fetal hematopoietic organ into the body’s central metabolic hub. However, how the liver acquires the capacity to adapt to carbohydrate nutrition, and how this process supports other organs, particularly the brain, remains poorly understood.

In infancy, liver function is immature and highly vulnerable to genetic or environmental insults. Dysfunction at this stage disrupts metabolism, causing hypoglycemia, hyperammonemia and energy failure, which in turn leads to neurological impairments that are often more severe than those seen in adulthood^4,5^. Defective liver development and function are hallmarks of pediatric metabolic disorders such as urea cycle defects, fatty acid oxidation disorders, and neonatal cholestasis, which often manifest as acute metabolic crises with lasting neurological consequences^6^. The post-weaning shift to carbohydrate-based nutrition therefore imposes unique demands on liver-brain metabolic coordination.

The liver-brain axis is a bidirectional network of circulating metabolites, cytokines, hormones, and neural signals^7^. Liver-derived nutrients, including amino acids, glucose, ketone bodies, and lipids, fuel the brain’s high energy demands and support neurotransmitter synthesis, synaptic plasticity, and myelination. Conversely, hypothalamic and brainstem circuits regulate hepatic glucose and lipid metabolism^8,9^. This interorgan communication maintains systemic metabolic homeostasis and enables adaptation to dietary and developmental cues. Disruption of this axis in early life can trigger both metabolic and neurodevelopmental defects.

Here, we define how the liver and brain coordinate their metabolic programs to adapt to the postnatal carbohydrate transition. We show that LACTB2 maintains mitochondrial RNA homeostasis to enable efficient translation of mtDNA-encoded ETC subunits during hepatic maturation, essential for developing oxidative capacity, glucose utilization, and detoxification. These findings reveal a fundamental mechanism by which dietary adaptation drives liver-brain axis maturation, establishing a low-toxicity, metabolically competent foundation for postnatal neurodevelopment.

## RESULTS

### Organ-specific metabolic reprogramming after birth

Mammalian postnatal life requires dynamic shifts in nutrient utilization, most notably the transition to a carbohydrate-rich diet after weaning. To investigate how metabolic programs coordinate this process, we performed transcriptomic and metabolomic profiling of the liver and brain (cerebral cortex) across six stages from birth to young adulthood (P0-P63; Fig. 1a), and used maSigPro^10,11^ to model organ-specific trajectories and identify time-resolved metabolic signatures. Both tissues exhibited marked declines in metabolites during the neonatal period (P0-P14; Fig. 1b,c), accompanied by progressive increases in a subset of metabolites (cluster 2). After weaning, the trajectories diverged, with liver metabolites continuing to rise (P28-P63), consistent with ongoing acquisition of metabolic capacity, whereas brain metabolites stabilized, reflecting the need for homeostatic balance during neuronal maturation.

**Fig. 1.**
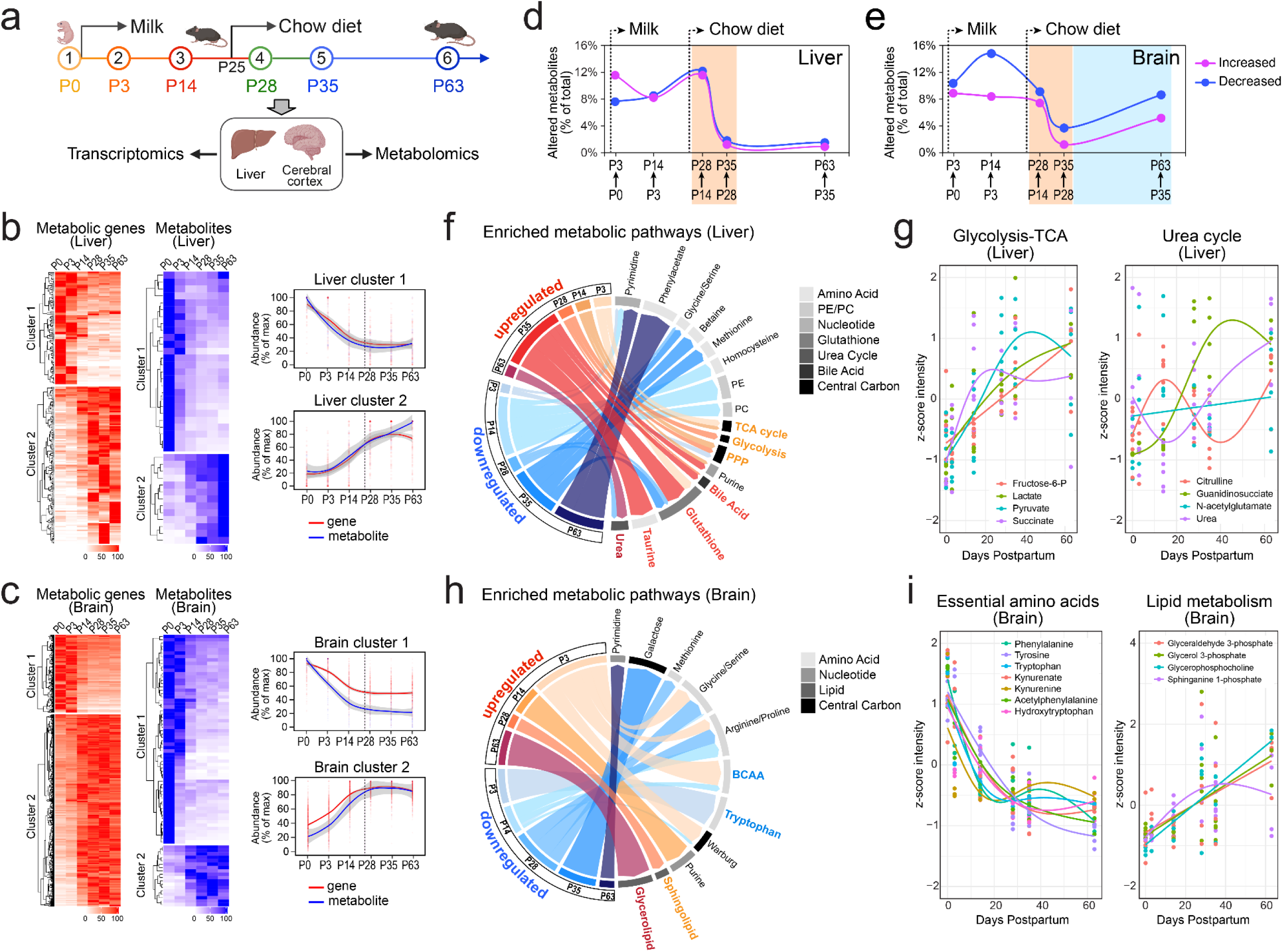
Liver-brain metabolic reprogramming during postnatal development. a. Schematic timeline of liver and brain collection across six postnatal stages, created with BioRender. b-c. Heatmaps of differential metabolic genes and metabolites during postnatal liver and brain development as analyzed by RNA-seq (*N* = 3-4 per stage) or metabolomics (*N* = 4-7 per stage). Genes and metabolites were grouped into two clusters by maSigPro and hierarchical clustering, with mRNA expression and relative metabolite levels shown as percentages of maximum abundance. Line graphs show the mean expression or metabolite abundance within each cluster. The lines are fitted curves with Loess regression analysis and shaded areas indicating 99% confidence interval. d-e. Stage-wise comparisons of differential metabolites during postnatal liver and brain development (fold change cutoff 1.5; *P* < 0.05). Pink indicates upregulation and blue indicates downregulation between consecutive stages, and shaded areas mark the metabolic transition phases. f. Circos plot showing temporal pathway enrichment of up- or down-regulated metabolites in the liver based on stage-wise comparisons in d. g. Z-score viewer of representative metabolite changes in glycolysis, the TCA cycle, and the urea cycle during postnatal liver development. h. Circos plot showing temporal pathway enrichment of up- or down-regulated metabolites in the brain based on stage-wise comparisons in e. i. Z-score viewer of representative metabolite changes in aromatic essential amino acid degradation and lipid metabolism during postnatal brain development.

These metabolic trajectories were accompanied by distinct, organ-specific gene expression changes. In the liver, upregulated genes were enriched for fatty acid oxidation, xenobiotic metabolism and cytochrome P450 pathways, while downregulated genes were associated with transcriptional regulation and epigenetic modification (Extended Data Fig. 1a,b). During this period, the brain upregulated neuronal signaling and synaptic pathways, while downregulating genes involved in RNA processing and regulation (Extended Data Fig. 1c,d). Thus, both tissues undergo transcriptional reprogramming, with the liver acquiring metabolic and detoxification capacity, and the brain specializing in neuronal function.

Because trajectory fits capture continuous patterns, we next performed stage-wise comparisons to pinpoint discrete transitions and pathway onsets. This analysis revealed rapid and pronounced metabolic shifts following the switch to chow diet in both organs (orange shading, Fig. 1d,e). In the liver, glycolysis, the TCA cycle, and the pentose phosphate pathway (PPP) were induced at P28, followed by increases in bile acids, taurine, and glutathione biosynthesis at P35, and strong induction of the urea cycle by P63 (Fig. 1f,g). In the brain, we observed substantial depletion of tryptophan, branched-chain amino acids (BCAAs), and their neurotoxic catabolites before weaning (P0-P14) (Fig.1h,i and Extended Data Fig. 1e), consistent with active clearance to establish early metabolic homeostasis. After weaning, brain metabolism stabilized by P35, and then entered a second transition (blue shading, Fig. 1e) marked by elevated metabolites for glycerolipid and sphingolipid biosynthesis, essential for membrane biogenesis and myelination^12^, and increased metabolites involved in neurotransmitter production and regulation (Fig. 1h,i and Extended Data Fig. 1f). Together, these analyses indicate that the liver and brain undergo temporally aligned but functionally distinct metabolic reprogramming, with the post-weaning carbohydrate transition severing as a key trigger for hepatic metabolic maturation and subsequent neural development.

### Nuclear-mitochondrial control of hepatic mitochondrial maturation

Liver maturation followed a defined developmental trajectory marked by progressive induction of metabolic pathways within the mitochondria, as assessed by metabolomics (Extended Data Fig. 2a,b). Gene set enrichment analysis revealed strong upregulation of nuclear-encoded mitochondrial genes, including several ETC components, while epigenetic regulators such as Polycomb Repressive Complex 2 (PRC2) and DNA methyltransferases, typically active during embryogenesis, were downregulated (Fig. 2a and Extended Data Fig. 2c). Hepatic mitochondria also underwent marked ultrastructural changes across postnatal stages. Electron microscopy showed sparse cristae at birth, partially electron-dense mitochondria by P14, and mature cristae-rich structures by P35 (Fig. 2b). Consistently, from P35 onward we observed significant increases in in oxygen consumption through complexes I and IV in isolated liver mitochondria, whereas complex II activity was unchanged (Fig. 2c). As mtDNA encodes 13 ETC subunits but excludes complex II, and mtDNA content remained stable across postnatal stages (Extended Data Fig. 2d,e), the rise in ETC activity reflects functional upregulation rather than increased mitochondrial mass.

**Fig. 2.**
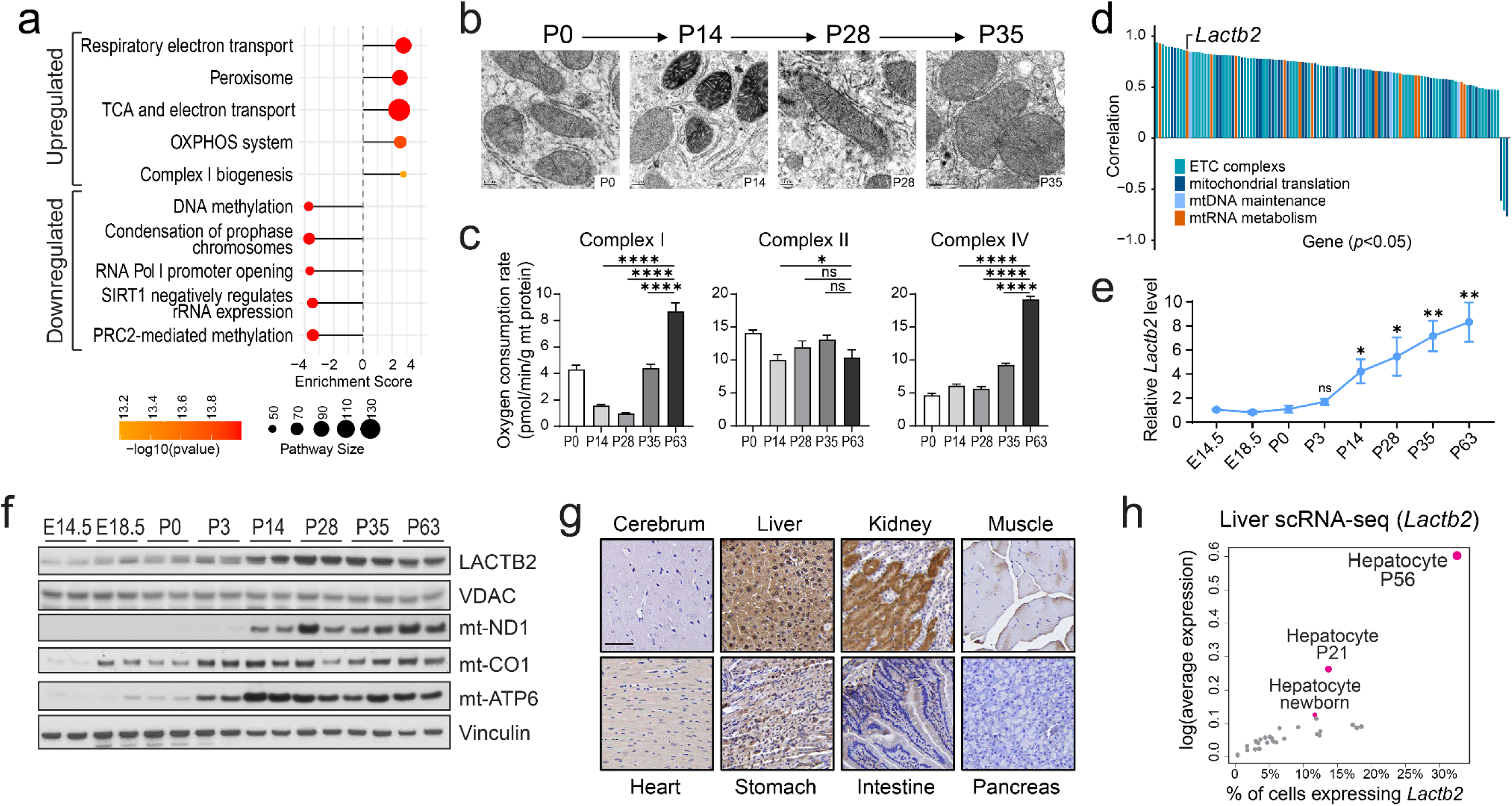
Co-induction of ETC components and LACTB2 during postnatal liver development. a. Top-ranked functional groups of differential genes significantly associated with developmental progression. b. Transmission electron microscopy (TEM) images of hepatic mitochondria across postnatal stages. Scale bar, 0.2 um. c. Comparison of ETC complex activity measured as oxygen consumption rate in isolated liver mitochondria across postnatal stages (*N* = 4-6 per group). Data are means ± SEM and analyzed by two-tailed unpaired t-test, \**P* < 0.05, \*\*\*\**P* < 0.0001. d. Waterfall plot of nuclear-encoded mitochondrial genes involved in mtDNA gene regulation and ETC function, ranked by their correlation with developmental progression (*P* < 0.05). e. Quantitative RT-PCR of *Lactb2* expression in developing livers (mean ± SEM, *N* = 4 per stage). Two-tailed unpaired t-test compared of each stage to P0. \**P* < 0.05, \*\**P* < 0.01. f. Immunoblot analysis of LACTB2, ETC complex proteins and mitochondrial VDAC in developing livers with vinculin as loading control. g. Immunohistochemical (IHC) staining of LACTB2 in major mouse organs using mouse tissue microarrays. Scale bar, 50 µm. h. scRNA-seq scatter plot showing *Lactb2* expression levels across distinct hepatic cell populations during postnatal development.

To define how hepatic mitochondrial maturation is regulated, we analyzed differentially expressed nuclear-encoded mitochondrial genes by function^13^. Approximately 40% were associated with oxidative phosphorylation, including structural components of the ETC and regulators of the mitochondrial central dogma, such as those controlling mtDNA transcription and translation (Extended Data Fig. 2f). Among these, *Lactb2*, which encodes a mitochondria-localized RNA endoribonuclease^14^, showed strong correlation with liver developmental progression (Fig. 2d), with transcript levels progressively upregulated from the perinatal period into adulthood (Fig. 2e). LACTB2 protein levels increased from P14 onward, coinciding with upregulation of mtDNA-encoded subunits such as ND1, CO1, and ATP6 (Fig. 2f). Immunofluorescence confirmed mitochondrial localization of LACTB2 (Extended Data Fig. 2g), with high expression specific to hepatocytes and kidney proximal tubules and absent in most other tissues (Fig. 2g). Single-cell RNA-seq further demonstrated hepatocyte-specific expression of *Lactb2* among >30 liver cell types during postnatal development^15^ (Extended Data Fig. 2h), with expression markedly increased by P21 and peaking in mature hepatocytes at P56 (Fig. 2h). Together, these results indicate that postnatal liver development involves the post-weaning maturation of oxidative phosphorylation through induction of both nuclear- and mtDNA-encoded ETC subunits, along with hepatocyte-specific upregulation of LACTB2.

### LACTB2 maintains hepatic mt-RNA homeostasis

LACTB2 is a 3’-5’ single-stranded RNA (ssRNA) endoribonuclease localized in the mitochondrial matrix^14^. It shares the catalytic core domain with ElaC Ribonuclease Z (ELAC) subfamily, including ELAC2, the only other known mammalian mitochondrial 3’-5’ ssRNA endoribonuclease required for mitochondrial tRNA (mt-tRNA) maturation (Extended Data Fig. 3a). However, the biological role of LACTB2 in mitochondrial function has remained undefined. To address this, we first performed metabolomic profiling in LACTB2-depleted H2.35 hepatocytes and identified marked metabolic changes associated with the TCA cycle (Extended Data Fig. 3b,c). LACTB2 depletion also caused a significant reduction in oxygen consumption, and re-expression of LACTB2 fully restored respiration (Extended Data Fig. 3d).

To directly assess mitochondrial RNA, we performed Northern blot analysis with strand-specific probes. Loss of LACTB2 caused accumulation of mature mt-mRNAs and mt-tRNAs, and this was reversed by LACTB2 re-expression (Fig. 3a). mt-rRNA abundance was not affected by LACTB2 loss. Functionally, LACTB2 depletion impaired assembly of all ETC complexes except complex II, which contains no subunits encoded by mtDNA, and reduced the activities of complexes I and IV (Fig. 3b,c). To determine whether accumulated mt-RNAs affect translation, we blocked mitochondrial protein synthesis with tigecycline^16^ and monitored protein recovery over time after drug withdrawal. LACTB2-depleted hepatocytes showed reduced expression of mtDNA-encoded ETC proteins, but not nuclear-encoded subunits such as SDHA or UQCRC2, and their recovery was markedly delayed (Fig. 3d). Therefore, with tissue-specific expression, LACTB2 adds a critical layer of mitochondrial RNA surveillance that preserves RNA homeostasis and supports efficient translation of mtDNA-encoded ETC subunits, particularly in hepatocytes.

**Fig. 3.**
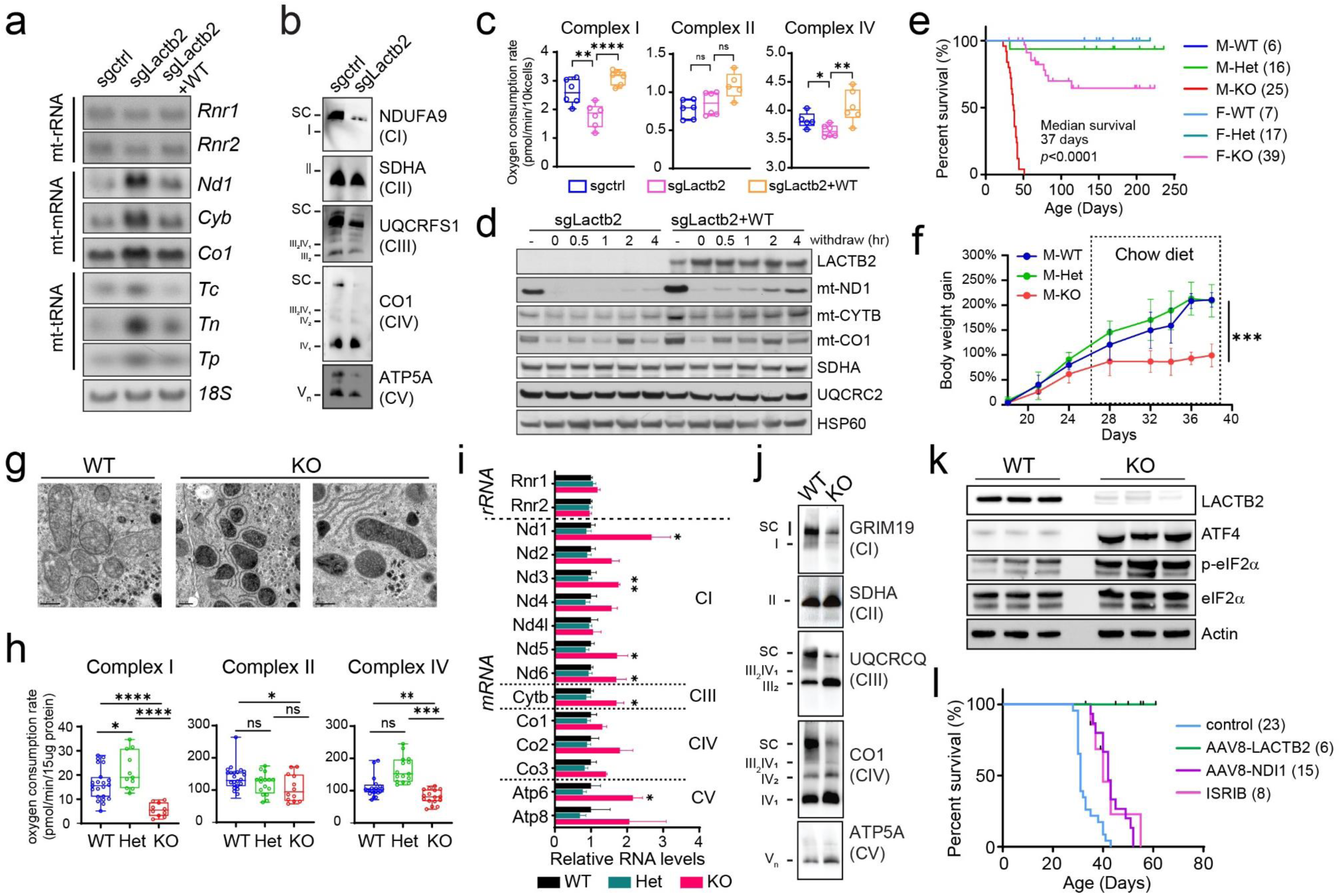
LACTB2 regulates mt-RNA homeostasis to support hepatic ETC function. a. Northern blot analysis of mt-RNAs in the indicated isogenic H2.35 hepatocytes. Nuclear 18S rRNA is used as a loading control. b. Blue-Native PAGE analysis of ETC supercomplexes in mitochondria isolated from control and LACTB2-depleted H2.35 hepatocytes. c. ETC complex activity assessed by oxygen consumption rates in the presence of complex-specific substrates in the indicated H2.35 hepatocytes. Results are means ± SEM (*N* = 6) and p values are calculated by two-tailed unpaired t-test, \**P* < 0.05, \*\**P* < 0.0, \*\*\*\**P* < 0.0001, ns, not significant. d. Immunoblot analysis of ETC complex proteins in LACTB2-depleted hepatocytes with or without LACTB2 re-expression. The cells were treated with tigecycline at 50 uM for 72h, followed by drug withdraw for the indicated hours. “-” indicates DMSO treatment. e. Survival of WT, Het and KO male and female mice. *N* numbers are indicated for each group; *P* value by log-rank Mantel-Cox test. f. Body weight gain up to 6 weeks postpartum. Data are mean ± SD (*N* = 4-6 per group) and \*\*\**P* < 0.001 by mixed-effects model. g. TEM of WT and KO male livers at P35 (*N* = 3 per group). Scale bar: 0.5 µm. h. ETC complex activity measured by OCR in isolated liver mitochondria from WT, Het and KO males. Results are mean ± SEM (*N* = 3-4 mice per group) and analyzed by two-tailed unpaired t-test. \**P* < 0.05, \*\**P* < 0.01, \*\*\**P* < 0.001, \*\*\*\**P* < 0.0001, n.s. not significant. i. Relative abundance of mitochondrial rRNAs and mRNAs in WT, Het and KO male livers at P35 as assessed by strand-specific RNA-seq (*N* = 4 per group). Results are mean ± SD and p values are calculated by two-tailed unpaired t-test, \**P* < 0.05, \*\**P* < 0.01. j. Blue-Native PAGE analysis of ETC supercomplexes using mitochondria isolated from WT and KO male livers at P35. k. Immunoblot analysis of ISR markers in WT and KO male mouse livers at P35 (*N* = 3 per group). l. Survival of KO male mice treated with AAV8-LACTB2, AAV8-NDI1 and ISRIB. *N* numbers are indicated for each group. *P* values by log-rank Mantel-Cox test: *P* < 0.0001 for AAV8-LACTB2 and AAV8-NDI1; *P* = 0.0022 for ISRIB.

### Hepatic mitochondrial maturation requires LACTB2

To assess the in vivo role of LACTB2, we generated a *Lactb2* knockout (KO) mouse strain via CRISPR-mediated targeting of exon3 of the *Lactb2* gene (del59fs), causing frameshift and loss of protein expression (Extended Data Fig. 3e; see Methods). Progeny carrying heterozygous *Lactb2* KO alleles showed reduced LACTB2 expression but displayed normal growth and fertility, and inbred heterozygotes (Het) produced offspring at Mendelian ratios (Extended Data Fig.3e,f). In contrast, homozygous KO males failed to gain weight after weaning (P25) and rapidly deteriorated, with a medium survival of 37 days (Fig. 3e,f). KO females were less severely affected, and most survived to adulthood and remained fertile, but with reduced average litter size (Extended Data Fig. 3g).

To assess the metabolic consequences of LACTB2 loss, we examined KO male livers at P35. EM analysis revealed dark and condensed mitochondrial structure with significantly reduced ETC activity (Fig. 3g,h), resembling the immature mitochondria normally seen at P14 (Fig. 2b,c). Strand-specific RNA-seq confirmed accumulation of mt-mRNAs, particularly complex I transcripts (Fig. 3i). Immunoblotting showed decreased protein abundances in complexes I, III, and IV, along with defective supercomplex assembly, while complex II and V were less affected (Fig. 3j and Extended Data Fig. 3h). Transcriptomic profiling further revealed that KO male livers at P35 deviated from normal developmental stages (Extended Data Fig. 3i). Unlike WT and Het samples, which clustered with mature stages, KO livers repressed adult maturation gene set and retained immature signatures (Extended Data Fig. 3j). Thus, LACTB2 is essential for hepatic ETC integrity and postnatal liver maturation, with male-biased vulnerability leading to premature lethality after weaning.

As complex I deficiency can induce the mitochondrial integrated stress response (ISR)^17^, we observed strong ISR activation in LACTB2-deficient male livers, marked by elevated p-eIF2α and ATF4 (Fig. 3k). To test whether restoring mitochondrial function could reverse these defects, we employed three strategies (Fig. 3l). First, KO males received a single injection of AAV8-LACTB2 at weaning were fully rescued from lethality, confirming that the phenotype stems primarily from hepatic loss of LACTB2. Second, liver-specific expression of yeast NADH dehydrogenase (NDI1), which bypasses complex I by transferring electrons directly to ubiquinone and regenerating NAD⁺^18^, partially rescued lethality. Third, treatment with the ISR inhibitor (ISRIB)^19^ from weaning also significantly improved survival. Together, these interventions demonstrate that restoring mitochondrial NAD⁺ or suppressing ISR activation can mitigate the effects of LACTB2 deficiency, suggesting that the acquisition of hepatic mitochondrial function is essential for postnatal adaptation to the carbohydrate transition.

### Liver immaturity disrupts brain metabolism after weaning

To evaluate systemic organ defects in KO males, we performed necropsy of 27 major organs with routine blood and chemistry tests. No gross nor any microscopic morphological abnormalities except rare liver lipidosis were observed, however KO males displayed hypoglycemia and elevated alkaline phosphatase (ALP) (Extended Data Fig. 4a). In vivo ^13^C-glucose tracing confirmed markedly reduced ^13^C labeling of glycolytic and TCA intermediates in the KO male liver, consistent with impaired carbohydrate utilization (Fig. 4a). KO female livers showed normal glucose metabolism (Extended Data Fig. 4b).

**Fig. 4.**
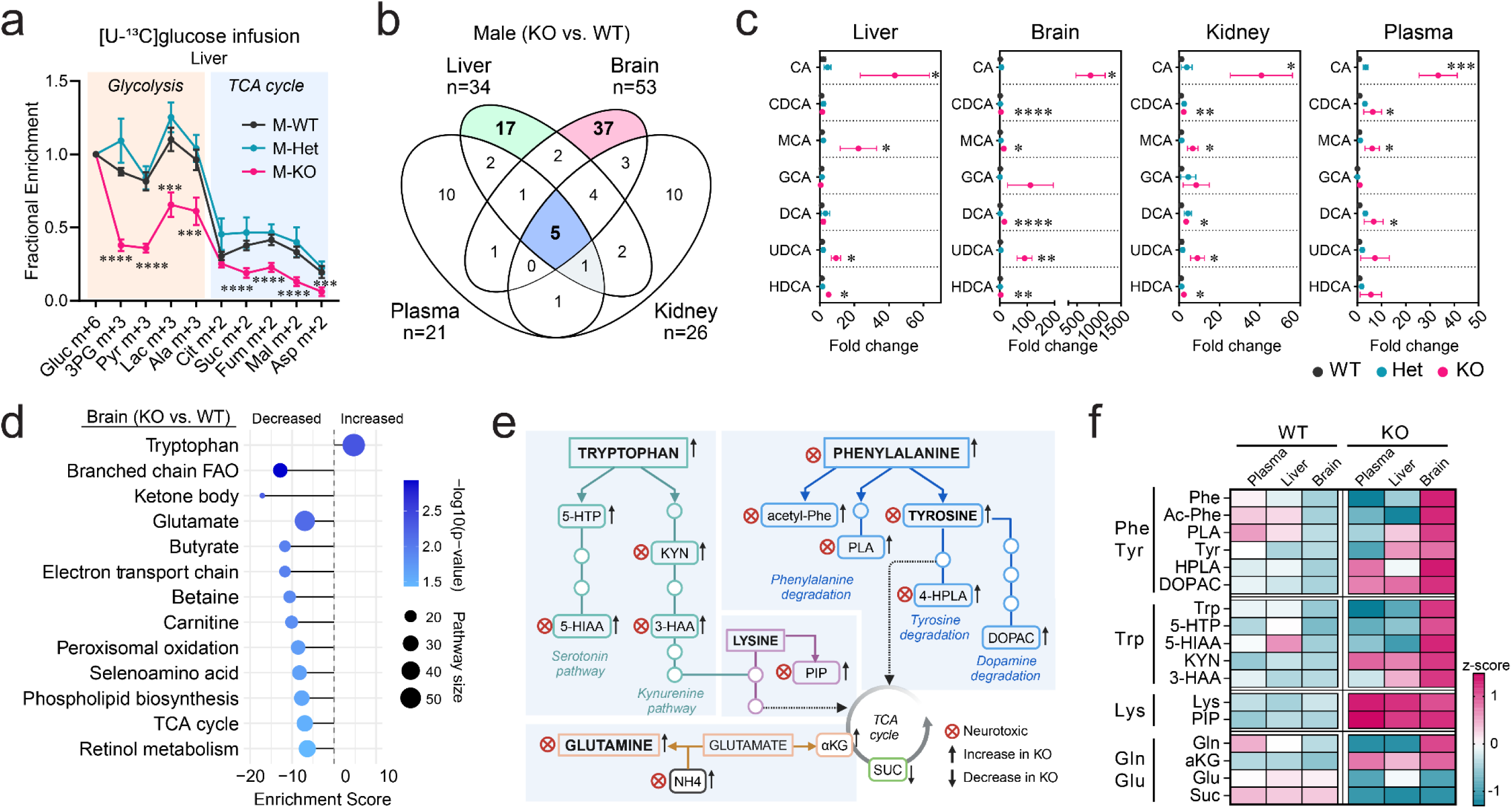
Loss of LACTB2 impairs glucose metabolism and induces neurotoxicity. a. In vivo [U-^13^C]glucose tracing showing ^13^C incorporation into glycolytic and the TCA cycle intermediates in male livers. Data are shown as mean ± SEM (*N* = 3 or 4 per group) with *P* values by two-tailed unpaired t-test, \*\*\**P* < 0.001, \*\*\*\**P* < 0.001. b. Venn diagram showing shared and tissue-specific differential metabolites identified in plasma, liver, brain and kidney based on metabolomic comparisons between WT and KO male mice at P35 (fold change cutoff 1.5; *P* < 0.05). c. Comparison of bile acid species in Het and KO livers relative to WT using LC-MS (*N* = 3-6 per group). Data are mean ± SEM and analyzed by two-tailed unpaired t-test, \**P* < 0.05, \*\**P* < 0.01, \*\*\**P* < 0.001, \*\*\*\**P* < 0.0001. Abbreviations: CA, cholic acid (primary); CDCA, chenodeoxycholic acid (primary); MCA, muricholic acid (rodent primary); GCA, glycocholic acid (primary conjugated); DCA, deoxycholic acid (secondary); UDCA, ursodeoxycholic acid (secondary); HDCA, hyodeoxycholic acid (secondary). d. Top enriched metabolic pathways with significantly altered brain metabolites between WT and KO males at P35. e. Schematic of amino acid catabolic pathways significantly altered in KO male brains, with accumulated neurotoxic intermediates highlighted. Abbreviations: 5-HTP, 5-hydroxytryptophan; KYN, kynurenine; 5-HIAA, 5-hydroxyindoleacetic acid; 3-HAA, 3-hydroxyanthranilic acid; PLA, phenyllactic acid; 4-HPLA, 4-hydroxyphenyllactic acid; DOPAC, 3,4-dihydroxyphenylacetic acid; PIP, pipecolic acid; αKG, α-ketoglutarate; SUC, succinate. f. Heatmap of relative metabolite abundance in dysregulated amino acid metabolism pathways in KO male brains, corresponding to pathways shown in e.

Transcriptomic analysis revealed sex-specific gene expression in the liver at P35. Females showed preferentially upregulated detoxification genes, particularly those involved in modification of bile acids and steroid hormones, while males biased toward higher lipid and glucose metabolism (Extended Data Fig. 4c,d). Integrated transcriptomic-metabolomic analyses confirmed stronger detoxification capacity in female livers, via xenobiotics and cytochrome P450 metabolism pathways, in contrast to enhanced bile acid synthesis and amino acid metabolism in male livers (Extended Data Fig. 4e-g). Consistent with this, KO males accumulated toxic metabolites, including liver bile acids and elevated blood ammonium (Extended Data Fig. 5a,b). Treatment with the ammonia scavenger sodium phenylbutyrate (120 mg/kg daily) significantly prolonged KO male survival (Extended Data Fig. 5c). When challenged with one high-dose carbon tetrachloride (CCl4) at weaning (5 ml/kg), KO males exhibited hepatocellular ballooning and necroptosis marker accumulation (p-MLKL and p-RIP3) within a week, further indicating immature detoxification and hepatocyte death (Extended Data Fig. 5d,e). Therefore, LACTB2 loss impaired hepatic adaptation to carbohydrate diet, leading to systemic metabolic toxicity, with males more vulnerable due to limited detoxification capacity during the development.

Given the liver’s central role in whole-body metabolism, we next assessed how LACTB2 loss impacts extrahepatic metabolism. Among liver, brain, kidney, and plasma, the most extensive changes occurred in the brain (Fig. 4b). Across all samples, bile acids and derivatives were consistently elevated, including highly toxic primary bile acids normally synthesized in the liver and modified by gut bacteria (Fig. 4c). Strikingly, KO male brains exhibited >100-fold increases in certain bile acid species relative to WT, far exceeding changes observed in circulation. As unconjugated bile acids, including hydrophobic cholic acid (CA) and deoxycholic acid (DCA), can cross the blood-brain barrier, their accumulation in KO male brains is likely neurotoxic, disrupting both metabolic homeostasis and neuronal signaling^20,21^.

Brain energy metabolism pathways were also disrupted, as evidenced by reduced ketone body and TCA cycle metabolites (Fig. 4d). Catabolites of aromatic essential amino acids, phenylalanine, tyrosine, and tryptophan, accumulated to high levels, particularly several neurotoxic metabolites linked to human inborn metabolic disorders^22^ (Fig. 4e). Some metabolites, such as α-ketoglutarate, lysine and its catabolite pipecolic acid accumulated systemically in plasma, liver and brain (Fig. 4f). By contrast, several others were specifically elevated in the brain despite relatively normal plasma and liver levels, indicating impaired local amino acid degradation. Thus, liver dysfunction caused by LACTB2 loss induced systemic metabolic toxicity and disrupted brain metabolic homeostasis, which recapitulated key features of inborn errors of metabolism with hepatic and neurological manifestations^23^.

### Postnatal brain myelination is susceptible to liver dysfunction

Perturbed brain metabolism and systemic toxicity during early childhood contribute to neurological defects and cognitive impairment^23,24^. Transcriptomic analysis of KO male brains revealed broad repression of neuronal development genes (Fig. 5a), many overlapping with gene loci linked to human neurodevelopmental disorders, manifesting seizures, developmental delay, and white matter anomalies according to OMIM clinical synopsis^25^ (Fig. 5b and Extended Data Fig. 6a). Consistent with these features, magnetic resonance imaging (MRI) of KO males showed increased total brain volume with disproportionate enlargement of the corpus callosum, a white matter tract essential for interhemispheric communication (Fig. 5c and Extended Data Fig. 6b). In WT mice, the corpus callosum is densely packed with myelinated axons, visualized by Luxol fast blue staining as organized, directionally aligned sheaths (Fig. 5d). By contrast, KO males exhibited profoundly disorganized myelin architecture, with TEM revealing broad abnormalities, including split myelin layers and ballooned myelin sheaths (Fig. 5e).

**Fig. 5.**
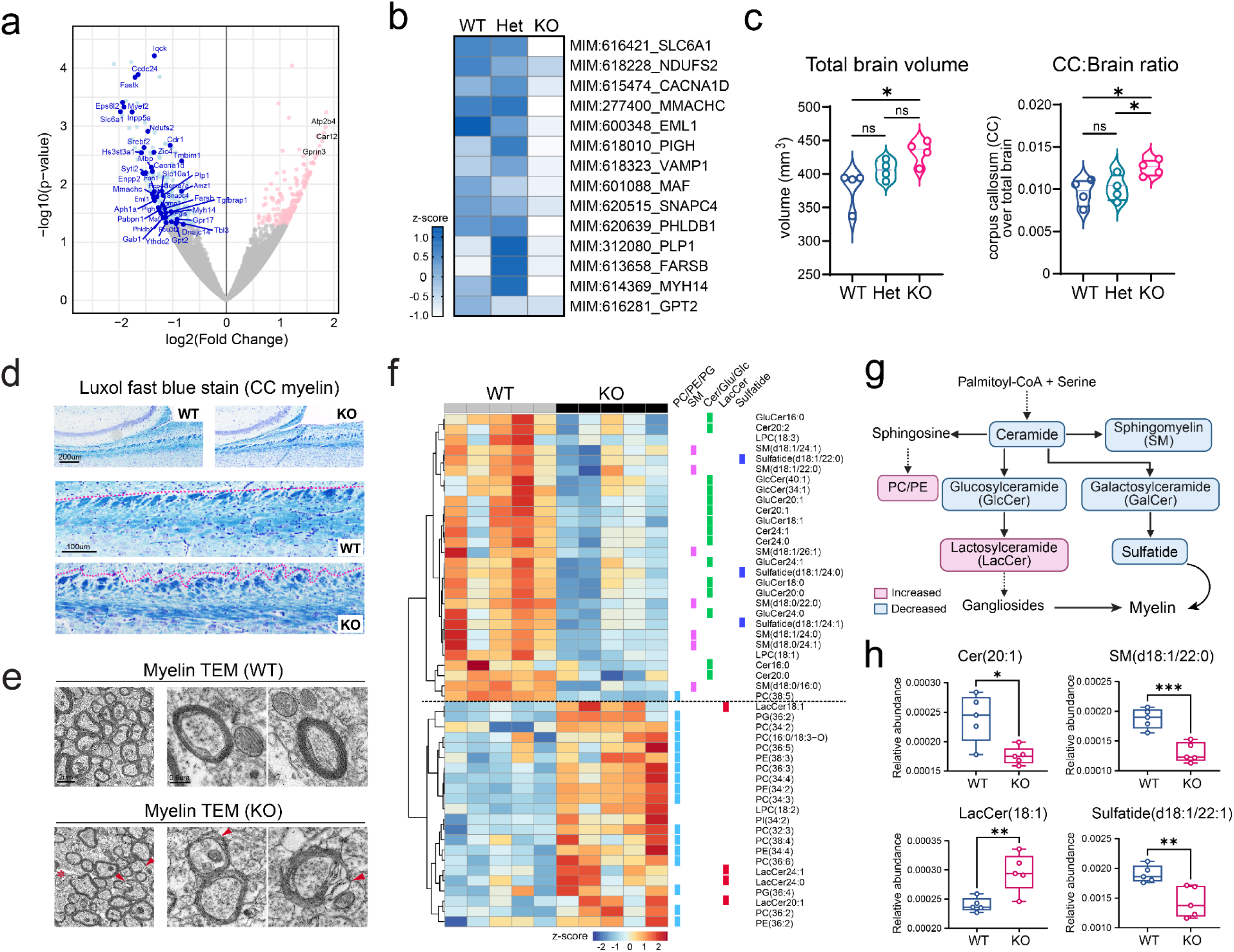
Corpus callosum defects in LACTB2 deficiency. a. Volcano plot of differential genes in the cerebrum between WT and KO males at P35 based on RNA-seq analysis (*N* = 3-4 mice per group). The genes involved in neuronal development and function are highlighted in blue or pink to indicate down- or up-regulation in KO vs. WT males. b. OMIM disease annotation of significantly downregulated genes in KO male brains that are known to cause human neurodevelopmental disorders. c. MRI quantification of total brain volume and corpus callosum (CC) fraction in WT, Het and KO male mice at P35 (*N* = 4 per group). Data are mean ± SD; two-tailed unpaired t-test, \**P* < 0.05; ns, not significant. d. Luxol Fast Blue staining of the corpus callosum showing distinct differences in myelin sheath integrity between WT and KO male brains at P35. e. TEM analysis of corpus callosum myelin in WT and KO male brains at P35. Red arrowheads indicate disrupted myelin layers; asterisks denote myelin sheath ballooning. Scale bars, 2 µm and 0.6 µm. f. Heatmap showing relative abundance of myelination-associated lipids in the corpus callosum of WT and KO male brains at P35. Lipid classes are color-coded: phospholipids (PC/PE/PG, blue), sphingomyelin (SM, pink), ceramides and derivatives (green), lactosylceramides (LacCer, red), and sulfatides (dark blue). g. Schematic of the myelin synthesis pathway with lipid species altered in the corpus callosum of KO male mice highlighted. h. Comparison of representative lipids in the corpus callosum between KO and WT males (*N* = 5 per group). *P* values were calculated by two-tailed unpaired t-test: \**P* < 0.05, \*\**P* < 0.01, \*\*\**P* < 0.001.

In mice, corpus callosum myelination begins around P10, accelerates at P17-P30, and continues to increase through three months of age^26,27^. Before weaning, thinner and immature sheaths are typical of early development (P14; Extended Data Fig. 6c). However, during the critical post-weaning window (P25-P35), when myelin expansion and compaction normally proceed rapidly^26^, KO males failed to establish proper sheath organization (Fig. 5d,e). Normal myelin maturation is characterized by shifts in phospholipid composition and enrichment of monounsaturated fatty acids (MUFA) in phospholipids and sphingolipids^28^, key structural components required for compact myelin assembly and signaling. To assess whether altered cerebral metabolism affects myelin lipid composition, we performed lipidomics on dissected corpus callosum. KO males showed persistently higher phosphoglycerolipids, accumulation of lactosylceramides (LacCer), and marked depletion of sphingomyelins and sulfatides (Fig. 5f-h). Because sphingomyelins and sulfatides are essential for forming the compact and insulated myelin sheaths that enables efficient nerve signal transmission, this imbalance in KO males disrupted the molecular foundation of myelin assembly during a critical period of postnatal brain maturation.

## DISCUSSION

Our study establishes hepatic mitochondrial maturation as a critical determinant of postnatal adaptation to a carbohydrate-rich diet and systemic metabolic stability. We identify LACTB2, a mitochondrial RNA endoribonuclease specifically expressed in hepatocytes and upregulated after birth, as a key regulator of mitochondrial RNA surveillance. By preventing accumulation of processed mt-mRNAs and mt-tRNAs, LACTB2 ensures efficient translation of mtDNA-encoded ETC subunits in the liver, thereby supporting respiratory complex assembly and sustaining glucose oxidation. LACTB2 deficiency impairs hepatic mitochondrial maturation, triggering the mitochondrial integrated stress response (ISR), which can be alleviated by restoring ETC function or inhibiting ISR. These findings uncover a previously unrecognized layer of developmental regulation by coordinating nuclear and mitochondrial genomes.

Early postnatal life requires high metabolic flexibility as the body transitions from milk to carbohydrate-based nutrition. Our study shows that this dietary adaptation coordinates the liver-brain maturation through temporal and organ-specific metabolic reprogramming, following developmental trajectories tailored to each organ’s physiological demands. After weaning, the liver acquires oxidative and detoxification capacity, while the brain first stabilizes core metabolism and later enhances synthesis of lipids and neurotransmitters to support neuronal specialization. LACTB2 deficiency disrupts post-weaning liver maturation and function, destabilizing brain metabolism during a vulnerable window, and impairing lipid remodeling required for corpus callosum myelination. Moreover, the brain’s limited detoxification capacity renders it sensitive to toxic metabolites circulating from defective liver metabolism or arising from local amino acid catabolism. This temporal interdependence highlights that liver mitochondrial readiness for glucose metabolism precedes brain maturation, providing both energetic substrates and a low-toxicity environment essential for neurodevelopment.

Sex-specific differences in postnatal liver development may determine the consequences of LACTB2 deficiency. Female livers develop stronger detoxification capacity, providing protection against toxic metabolite accumulation, whereas males are more vulnerable to systemic toxicity and premature lethality. However, this protection in females may be context-dependent, and metabolic stresses such as pregnancy or lactation impose additional demands on detoxification and mitochondrial function^29,30^. This may explain the reduced average litter sizes observed in pregnant KO females. Given male lethality occurs near sexual maturation, these sex-dimorphic features of hepatic metabolism likely influence the trajectory of liver maturation and its resilience to stress, particularly during periods of active hormone changes such as puberty.

Our findings have broader implications for early-life health and disease risk. In humans, the prolonged course of brain development suggests that liver functional maturation may remain critical over a longer window than in mice^31^. The staged transition from milk to carbohydrate-rich foods not only supports digestive adaptation^32^ but also allows hepatic mitochondrial and detoxification pathways to mature. Disruption of this program by genetic defects or environmental insults can overwhelm the immature liver and trigger systemic metabolic crises. Pediatric disorders such as urea cycle defects^33^, fatty acid oxidation disorders^34^, and hepatic encephalopathy^35^ often manifest in early childhood with neurological impairments similar to those observed in LACTB2-deficient mice. Interventions that rescue these defects, such as ISR inhibition, NAD+ restoration, or liver-directed LACTB2 gene delivery, demonstrate that enhancing hepatic mitochondrial competence is a promising strategy for improving early-life liver health, with broad relevance to metabolic disorders and long-term vulnerabilities to metabolic and neurological dysfunction.

## ACKNOWLEDGEMENTS

We thank the following shared resources at St. Jude Children’s Research Hospital: Comparative Pathology Core, Cell and Tissue Imaging Center, Vector Development and Production Laboratory, and Center for In Vivo Imaging and Therapeutics (CIVIT), supported by NCI (P30 CA021765) to the St. Jude Comprehensive Cancer Center. We thank Dr. Hao Zhu and the mouse genome engineering facility at Children’s Research Institute of UTSW for mouse model generation, the Advanced Technology and Genomic (ATG) Core and the Center of Excellence for Leukemia Studies (CELS) at St. Jude for technical assistance, and Dr. SeungHye Han for advice on in vivo ISRIB administration. This work was supported by the American Lebanese Syrian Associated Charities (ALSAC) (to J.X. and M.N.), the National Institutes of Health (NIH) grants R01DK111430, R01CA230631, R01CA259581 (to J.X.), and R35GM151245 (to S.M.N.). R.J.D is a Howard Hughes Medical Institute (HHMI) Investigator and supported by NIH grant R35CA220449 and CPRIT grant RP240494. Z.W. is supported by NIH grant 1K99GM154116.

## CONTRIBUTIONS

M.N. conceived the study. Z.W., H.C., H.C., X.G., Y.J., C.P., H.C., C.G.R., J.D.S., S.M.N and M.N. performed experiments and analyzed the data. H.S.V generated and analyzed metabolomics and isotope tracing data. Y.Z., D.C., S.A. and M.N. performed bioinformatic analyses. P.N. performed protein structure analysis. L.J.J. assisted in necropsy analysis and report. J.X., R.J.D, and M.N. interpreted the results and wrote the manuscript.

## Methods

### Cell lines and cell culture

Human Huh7 liver tumor cells (a gift from Dr. Hao Zhu at UTSW) and mouse H2.35 hepatocytes (ATCC, CRL-1995) were cultured in DMEM (Sigma, D5796) supplemented with 5% heat-inactivated fetal bovine serum (FBS) at 37°C with 5% CO_2_. To deplete *Lactb2* in H2.35 cells, cells were transfected with the PX458 construct that contains sgRNA against mouse *Lactb2* gene, followed by sorting of highest 5% GFP signal using flow cytometer (BD FACSAria II).

### Mice

All animal procedures were approved by the Institutional Animal Care and Use Committees at UT Southwestern Medical Center and St. Jude Children’s Research Hospital in accordance with the Guide for the Care and Use of Laboratory Animals. C57BL/6J mice were purchased from the Jackson Laboratory (Strain #000664). *Lactb2* constitutive knockout (KO) mice were generated by the Mouse Genome Engineering Core of the Children’s Research Institute at UTSW using CRISPR/Cas9 genome editing technology. Guide RNA was designed to target the sequence in exon 3 of *Lactb2* gene. Guide RNA, S. pyogenes Cas9 mRNA, and a single-strand oligo donor containing mutations were injected into single celled zygotes from C57BL/6J mice. C57BL/6J WT females were crossed to *Lactb2* KO founder males to confirm germline transmission. Interbreeding of *Lactb2^+/-^* male and female littermates was performed to obtain homozygous *Lactb2^-/-^* mice. All mice were housed in a pathogen free environment with a 12-hour light-dark cycle, 75°F and 35% humidity.

Mouse necropsy was performed by the St. Jude Comparative Pathology Core, with comprehensive pathological and histological analyses conducted on 27 tissues, including the heart, skeletal muscle, tongue, thyroid glands, lungs, esophagus, liver, kidneys, adrenal glands, thymus, spleen, lymph nodes, salivary glands, stomach, small intestine, cecum, colon, pancreas, reproductive organs, skin, brain, pituitary gland, eyes, ears, Harderian glands, bone marrow, and vertebral column. Complete blood count (CBC), blood glucose, lactate, and ammonia levels, along with liver function tests, were also assessed. For ISR inhibition, mice received intraperitoneal injections of the ISR inhibitor ISRIB (2.5 mg/kg) every other day^17^ starting at weaning (P25). For ammonia detoxification, sodium phenylbutyrate (SPB) was administered intraperitoneally at 120 mg/kg daily^36^, also beginning at weaning.

### Plasmids

The coding regions of mouse *Lactb2* gene and yeast NDI1 gene were PCR amplified using the primer pairs listed in Supplemental Table 1. The PCR products were purified using Qiagen PCR purification kit and then subjected to NEBuilder HiFi DNA Assembly reaction with the linearized pAAV-MCS vector, following the manufacturer’s protocol. For lentiviral Lactb2 cloning, the PCR products were digested and inserted between AsiSI and MluI sites of pLenti-EF1a-Myc-DDK-IRES-Puro vector. The positively selected clones were confirmed by Sanger sequencing.

### RNA-seq analysis

Total RNA was isolated from mouse liver and brain tissues using miRNeasy mini kit (Qiagen, #217084), followed by rRNA depletion using NEBNext® rRNA Depletion Kit (NEB, E6350). RNA-seq library was prepared using the NEBNext® Ultra II Directional RNA Library Prep Kit for Illumina (NEB, E7760L) and applied to paired-end sequencing on Illumina NextSeq500 or NextSeq2000. Sequencing reads from all RNA-seq experiments were aligned to mm10 reference genome by STAR v. 2.5.2b. Output BAM files were converted to BED format using the “bamtobed” command from BEDtools v.2.30.0. BED files were then converted to a normalized wiggle file using a custom python script. Normalized wiggle files were then converted to bigwig format using wigToBigWig with “-clip” parameter. Genic read counts were derived using HTSeq with parameter “-s yes -- nonunique none” using the GTF file downloaded from UCSC. Raw read counts were normalized by DESeq2^37^.

Genes with dynamic temporal profiles (DDGs) were identified using maSigPro, an R package designed for transcriptomic time courses^10^. Normalized read count from DESeq2 were used as input, and genes with less than 10 reads in all time points were excluded. maSigPro was run with a degree=4 (polynomial), count=TRUE (count data), Q=0.01 and alfa=0.01. A k=2 was set to identify two clusters of genes with DDGs, considering a goodness-of-fit (R2) of at least 0.6. Heatmaps were generated using the R package “ComplexHeatmap”, within which the percentage (%) of the maximum value across all stages was calculated for each gene in the two clusters. We used hypergeometric distribution model to test CPDB pathways^38^ that are enriched with these gene clusters.

### Metabolomics

About 20 mg of tissues were homogenized in ice-cold 80% HPLC-grade methanol. After three freeze-thaw cycles in liquid nitrogen, the lysates were centrifuged at 12,000 rpm at 4°C for 15 min and the supernatants were collected and dried in a SpeedVac concentrator (Thermo Savant). Data acquisition was performed by reverse-phase chromatography on a 1290 UHPLC liquid chromatography (LC) system interfaced to a high-resolution mass spectrometry (HRMS) 6546 Q-TOF mass spectrometer (MS) (Agilent Technologies, CA). The MS was operated in both positive and negative (ESI+ and ESI-) modes. Analytes were separated on an Acquity UPLC® HSS T3 column (1.8 μm, 2.1 x 150 mm, Waters, MA). The column was kept at 40°C. Mobile phase A composition was 0.1% formic acid in water and mobile phase B composition was 0.1% formic acid in 100% ACN. The LC gradient was 0 min: 1% B; 5 min: 5% B; 15 min: 99%; 23 min: 99%; 24 min: 1%; 25 min: 1%. The flow rate was 250 μL min-1. The sample injection volume was 5 μL. ESI source conditions were set as follows: dry gas temperature 350°C and flow 12 L min-1, fragmentor voltage 175 V, sheath gas temperature 350 °C and flow 12 L min-1, nozzle voltage 500 V, and capillary voltage +3500 V in positive mode and −3500 V in negative. The instrument was set to acquire over the full m/z range of 40–1700 in both modes, with the MS acquisition rate of 1 spectrum s-1 in profile format.

Raw data files were processed using Profinder B.08.00 SP3 software (Agilent Technologies, CA) with an in-house database containing retention time and accurate mass information on 600 standards from Mass Spectrometry Metabolite Library (IROA Technologies, MA), which was created under the same analysis conditions. The in-house database matching parameters were: mass tolerance 10 ppm; retention time tolerance 0.5 min. Peak integration result was manually curated in Profinder for improved consistency. For some compounds, when standards were analyzed by the same experimental setup, more than one chromatographic peak with similar qualification but different intensity were detected most likely due to interaction with column and high pressure; each chromatographic peak is denoted by a suffix (-1 or -2 etc). The peak area for each detected metabolite was normalized against the total ion count (TIC) followed by statistical analyses and pathway enrichment analysis by Metaboanalyst 6.0^39^.

Similar to the time series analysis of RNA-seq data, we performed maSigPro analysis for metabolomics data with a degree=4 (polynomial), count=FALSE (is not count data), Q=0.01, and alfa=0.01. We set k=2 to identify two clusters of metabolites with DDGs. For each metabolite in the clusters, we calculated the percentage (%) of maximum value across all stages and used this value to draw heatmaps using the R package “ComplexHeatmap”.

### Lipidomics

Approximately 10 mg of corpus callosum was dissected on dry ice and placed into a 1.5 mL Eppendorf tube containing 250 µL of methanol and 200 µL of water. The tissue was homogenized on ice using a Fisherbrand 150 Handheld Homogenizer Motor (Thermo Fisher Scientific, MA, USA). Subsequently, 750 µL of cold methyl tert-butyl ether (MTBE) was added, and the mixture was vortexed for 10 s and centrifuged at 15,000 × g for 10 min. The upper MTBE layer was collected and dried using a SpeedVac. Dried samples were dissolved in 100 µL of methanol: toluene (9:1, v:v) and 5 µL was injected for mass spec analysis. Metabolites were separated using Zorbax EclipsePlus C18, 100 x 2.1 mm, 1.8 µm (Agilent Technologies, CA). The column was kept at 60°C. Mobile phase A is 60:40 (v:v) acetonitrile:water with 10 mM ammonium formate and 0.1 % formic acid. Mobile phase B is 90:10 (v:v) isopropanol:acetonitrile with 10 mM ammonium formate and 0.1 % formic acid. The gradient was 0 min: 15% B; 2 min: 30% B; 2.5 min: 48% B; 8.5 min: 72% B; 20 min: 99% B; 20.5 min: 99% B; 21 min: 15% B; 25 min: 15% B. Lipidomics data were processed in Profinder, as described for metabolomics, with raw data manually curated using a mass tolerance of 10 ppm and a retention time tolerance of 0.5 min.

### Quantitative real-time PCR

To evaluate mitochondrial DNA content, genomic DNA was isolated from 25 mg of mouse liver tissues with QuickExtract DNA extraction solution (Epicenter, QE09050) according to the manufacturer’s instruction. 100 ng of total genomic DNA was used to amplify *mt-Cytb* and nuclear gene *Rpl13a* using the SYBR Green mix (Bio-Rad). To examine *Latctb2* expression levels in developing mouse livers, 500 ng of total RNA was used for complementary DNA (cDNA) synthesis with the iScript^TM^ cDNA synthesis kit (Bio-Rad), followed by quantitative real-time PCR using the SYBR Green mix (Bio-Rad).

### Lentivirus transduction

Lentivirus was produced by transfection of HEK293T cells using the Lentiviral Packaging Kit (Origene, TR30037). Viral supernatants were harvested at 48 hours and 72 hours, filtered through a 0.45 µm filter and concentrated using PEG-it Virus Precipitation Solution (System Biosciences, LV810A). For transduction, the lentiviral pellets were suspended in culture medium and added to H2.35 hepatocytes at 70-80% confluency per well in 6-well plates. After 48 hr of transduction, the cells were selected under 0.35 µg/mL puromycin (Thermo Fisher, NC9138068) for 1-2 weeks for stable expression of LACTB2.

### Transmission electron microscopy (TEM)

For EM analysis of mouse liver mitochondria, anesthetized mice were transcardially perfused with fresh 4% paraformaldehyde and 1% glutaraldehyde in 0.1M sodium cacodylate buffer for approximately 7 minutes with volume and rate: 10.0 ml running at 1.25 mL/minute. Liver segments no larger than 1 mm^3^ were grossly trimmed from the major lobes of dissected livers following perfusion. Liver cubes obtained from grossing were further fixed by immersion in 2.5% (v/v) glutaraldehyde in 0.1M sodium cacodylate buffer prior to dehydration and embedding. The samples were then rinsed in 0.1M sodium cacodylate buffer and post-fixed in 1% osmium tetroxide and 0.8% potassium ferricyanide in 0.1M sodium cacodylate buffer for 1.5 hours at room temperature. After three rinses in water, they were en-bloc stained with 4% uranyl acetate in 50% ethanol for 2 hours, dehydrated with increasing concentrations of ethanol, transitioned into resin with propylene oxide, infiltrated with EMbed-812 resin and polymerized in a 60°C oven overnight. Blocks were sectioned with a diamond knife (Diatome) on a Leica Ultracut 7 ultramicrotome (Leica Microsystem) and collected onto copper grids, post-stained with 2% aqueous uranyl acetate and lead citrate. Images were acquired with a Tecnai G2 spirit transmission electron microscope (FEI, Hillsboro, OR) equipped with a LaB6 source at 120kV using a Gatan Ultrascan CCD camera.

For EM analysis of myeline in corpus callosum, mice were anesthetized and perfused with 3% glutaraldehyde at 1 mL/min for approximately 15 minutes. Brain was dissected and positioned dorsal side up in a brain matrix to isolate corpus callosum sections, as previously described^40^. Sections (0.5 mm thick) were collected from the same anatomical location across all mice and post-fixed in 3% glutaraldehyde at 4°C overnight. The corpus callosum sections were cut in half and embedded in 4% low melt agarose (Fischer). The samples were then rinsed in 0.1M sodium cacodylate buffer and post-fixed in 2% osmium tetroxide in 0.1M sodium cacodylate buffer for 1 hour at room temperature. They were then rinsed three times in sodium cacodylate, followed by three rinses in water, a rinse in 30% ethanol and an additional rinse in water. Then they were enbloc stained with 4% aqueous uranyl acetate 1 hour, dehydrated with increasing concentrations of ethanol, transitioned into resin with propylene oxide, infiltrated with EMbed-812 resin and polymerized in a 60°C oven for 48 hours. Blocks were sectioned with a diamond knife (Diatome) on a Leica Ultracut 7 ultramicrotome (Leica Microsystem) and collected onto copper grids. Images were acquired with a Tecnai F20 transmission electron microscope (FEI, Hillsboro, OR) using a NanoSprint15 sCMOS camera (AMT).

### Mitochondrial electron transport chain activity assay

Respiration rate in H2.35 hepatocytes or isolated liver mitochondria were determined by measuring oxygen consumption rate using Seahorse bioscience extracellular flux (XFe96) analyzer following the manufacturer’s instruction. Mouse livers (∼200-400 mg) were homogenized in 1 mL isolation buffer (5mM HEPES, 70mM sucrose, 220mM mannitol, 5mM MgCl2, 10mM KH2PO4, 1mM EGTA, pH 7.2) on ice. Supernatant containing mitochondria was collected after centrifuged at 600 xg for 5 minutes at 4°C, followed by a higher-speed centrifugation at 10,000 xg for 10 minutes to enrich mitochondria. After two washes with 1 mL isolation buffer, the mitochondrial pellets were re-suspended in isolation buffer and the protein quantification was determined using DC protein assay Kit II (Bio-Rad #5000112). For Seahorse assay, 20 µL mitochondrial samples with 15 µg proteins were loaded in each well. After incubating on ice for 10 min, the plate was centrifuged at the maximum speed for 2 minutes at 4°C. Immediately ETC complex-specific substrates were added to the cells, and the plate was subjected to the Seahorse cell mito stress test according to the manufacturer’s instruction. ETC complex activities of H2.35 cells were evaluated using the Seahorse XF Plasma Membrane Permeabilizer (PMP) system (Agilent, 102504-100), followed by measurement of specific substrate oxidation using the cell mito stress test. Substrates used for ETC complexes include: 10mM glutamate and 1 mM malate for complex I, 5 mM succinate and 2 µM rotenone for complex II, and 10 mM ascorbate, 100 µM TMPD and 2 µM antimycin A for complex IV.

### Immunofluorescence microscopy

Cells were seeded on coverslips that were coated with 10 µg/mL fibronectin (Sigma-Aldrich, F1141-5MG) in PBS for 1 hour at 37°C. After fixation with fresh warm 4% paraformaldehyde in PBS for 15 minutes at room temperature, cells were permeabilized in 0.1% (v/v) Triton X-100 in PBS for 10 minutes, pre-blocked with filtered 1% BSA (w/v) in PBS for 1 hour and then incubated with primary antibody against LACTB2 (1:200, Proteintech, 16783-1-AP), and HPS60 (1:100, ThermoFisher, MA3-012) diluted in the blocking buffer for 1 hour at room temperature. After 3x 5-min wash with PBS, coverslips were then incubated with secondary antibodies (1:500, Alexa Fluorophores 488 and 555) diluted in the blocking buffer for 1 hour in dark at room temperature. After 3x 5-min wash with PBS in dark, coverslips were mounted onto slides using Prolong Antifade Mountant (Invitrogen, P36935) overnight. Cells were imaged using Laser scanning confocal Zeiss LSM880 (x63 objective), with Z-stacks acquired. Images were processed using ImageJ software.

### Immunohistochemistry

Mouse liver tissues were collected and fixed in neutral buffered 10% formalin for 48 hours before submission to the histology core for preparation of FFPE blocks and slides. Immunohistochemical (IHC) staining was performed on FFPE slides and mouse tissue microarrays (Zyagen, MAP-MT2 and MAP-MT3) following the standard procedure as described previously^41^. The primary antibodies for IHC staining include anti-LACTB2 (1:500, Proteintech), anti-Phospho-MLKL(Ser345) (1:500, Cell Signaling Technology), and anti-Phospho-RIP3 (Thr231/Ser232) (1:500, Cell Signaling Technology).

### Luxol fast blue-cresyl violet staining of myelin

Deparaffinized cerebral slides were incubated overnight with the Luxol Fast Blue solution (Poly Scientific s235). Slides were rinsed in 95% alcohol, then distilled water before beginning the differentiation process via immersion in 0.05% lithium carbonate solution (Poly Scientific, s2048) for 10-20 seconds. Slides were then placed in 70% alcohol until gray and white matter could be distinguished. After rinsing in distilled water, slides were incubated in Cresyl Echt Violet solution (Poly Scientific, s167c) at 60° C for 6 minutes. After several changes of 95% alcohol, slides were dehydrated in absolute alcohol, cleared in xylene, and mounted with permount. Slides were analyzed using an Olympus BX53 Upright microscope (Olympus, Tokyo, Japan) and a NanoZoomer 2.0HT scanner (Hamamatsu Photonics, Hamamatsu City, Japan).

### Western blot

Whole cell lysates were extracted from mouse liver or hepatocytes in RIPA buffer (Boston Bioprodcuts, BP-115) followed by three freeze/thaw cycles. The protein supernatants were quantified using DC protein assay Kit II (Bio-Rad #5000112). Proteins were separated on NuPAGE 4-12% Bis-Tris protein gels (ThermoFisher, NP0323), transferred to PVDF membranes, and probed with primary antibodies and the corresponding secondary antibodies as listed in Supplementary Table 1. Immunoreactive proteins were visualized by enhanced chemiluminescence (Pierce, 32106).

### Northern blot

Total RNA was isolated from hepatocytes using miRNeasy mini kit (Qiagen, #217084) according to the manufacturer’s protocol. Northern blot analysis was performed as described previously^42^. Oligonucleotide probes were designed to be perfectly complementary to the mature mitochondrial rRNA, mRNA or tRNA sequences and Ultrahyb-oligo buffer (Ambion, #8663) was used for hybridization. The probe sequences were listed in Supplemental Table 1.

### Blue Native PAGE

Crude mitochondrial fractions from cells or liver tissues were isolated and Blue Native (BN) PAGE gel analysis was performed as previously described^43^. Briefly, 100 µg of mitochondria were pelleted at 10,000 x*g* for 10 min at 4°C and re-suspended in 1x pink lysis buffer (Invitrogen, BN20032) with addition of Digitonin (GoldBio) to a final concentration of 1% (w/v). After incubation on ice for 15 minutes, samples were spun at 20,000 g for 20 minutes. 6 µL of NativePAGE sample buffer (Invitrogen, BN20041) was added and 10 µL of sample was run on 3-12% NativePAGE gels (Invitrogen, BN2011B) with NativePAGE anode buffer and dark blue cathode buffer at 150 V for 1 hour then switched to light blue cathode buffer and run at 30 V overnight. After gel transfer, the membrane was incubated with the indicated primary antibodies, followed by the corresponding secondary HRP antibodies. SuperSignal West Femto Maximum Sensitivity Substrate (Thermo, 34096) was used to visualize protein bands.

### Adeno-Associated Virus (AAV) production

To prepare AAV particles carrying the coding regions of mouse *Lactb2* gene or yeast *NDI1* gene, SJ293TS-DPB cells were transiently transfected with AAV vector and helper plasmids, pCR+LTAAVhelp2-8 and pHGTI-Adeno1 at a plasmid ratio of 1:1:3, respectively, using TransIT-VirusGEN (Mirus Bio) at a ratio of reagent to DNA of 1.5:1. Forty-eight hours post-transfection, cells were collected by centrifugation at 930 x *g* for five minutes and suspended in TD buffer (140 mM NaCl, 5 mM KCl, 0.71 mM K_2_HPO_4_, 25 mM Tris pH 7.5 and 6.4 mM MgCl_2_). To lyse cells, Triton X-100 was added to a final concentration of 0.8%, and the cell suspension was subjected to three freeze-thaw cycles. The cell lysate was then treated with Benzonase (Millipore-Sigma) at 100 units per mL at 37°C for one hour to remove extracellular plasmid and genomic DNA. Sodium chloride was added to a final concentration of 1.15M, and the lysate was clarified by centrifugation at 2,500 x *g* for twenty minutes. Viral particles were purified using a discontinuous iodixanol density gradient via ultracentrifugation at 200,000 ×*g* at 18°C for one hour. The AAV containing fraction was filtered using a 0.22 mM PES filter and loaded onto a POROS™ GoPure™ HQ50 prepacked column (ThermoFisher Scientific) using an Akta Avant chromatography system (Cytiva) according to the manufacturer’s instructions. Viral particles were eluted using 20 mM Bis-Tris propane pH 9.0 with 1 M NaCl. Buffer exchange and concentration was performed using a 30 kDa Vivaspin centrifugal concentrator (Sartorius AG, Göttingen, Germany) by diluting eluate to a total volume of 15 mL using PBS and centrifuging at 3,000 × g at 18°C for thirty minutes to reduce volume to approximately 0.5 mL. This process was repeated a total of three times. The concentrated and purified AAV vector was formulated with Pluronic-F68 (Millipore-Sigma) to 0.02% final concentration, passed through a 0.22 mM PES filter, aliquoted and stored at -80°C. Titration of AAV vectors was performed using a modified ddPCR protocol^44^. Briefly, 10 uL of the vector preparation was treated with 10 units of DNaseI (NEB) at 37°C for thirty minutes and then, serially diluted 10-fold using GeneAmp™ 10x PCR buffer (Thermo Fisher Scientific) containing 0.05% Pluronic-F68 (Millipore-Sigma) and 2 ng per mL sheared salmon sperm DNA (Thermo Fisher Scientific). Five microliters of DNaseI treated and diluted vector from dilutions 10^-1^ to 10^-8^ were used as template in PCR using a QX200 digital droplet PCR system (Bio-Rad, Carlsbad, CA, USA) according to the manufacturer’s instructions. The following primer-probe set was used to amplify the AAV ITR sequence, ITR-F: 5’-GGAACCCCTAGTGATGGAGTT-3’, ITR-R: 5’- CGGCCTCAGTGAGCGA-3’ and ITR-Probe: 56- HEX/CACTCCCTC/ZEN/TCTGCGCGCTCG/3IABkFQ under the following conditions; 95°C for 10 minutes for one cycle; 40 cycles of 94°C for 30 seconds, ramp speed 2°C per second, 60°C for 60 seconds, ramp speed 2°C per second and one cycle of 98°C for 10 minutes and held at 4°C. Vector titer per mL was calculated by dividing the number of copies per reaction well by 5 × dilution factor. The viral particles were stored as aliquots in -80°C freezer. For AAV8 transduction of mouse liver, each mouse was injected with 5x10^10^ genome copies of AAV8 vector from retro-orbital plexus during weaning at P25.

### Infusion and stable isotope tracing

Mice were fasted overnight and then subjected to infusion procedures at 10am-12pm. After anesthesia with Ketamine/Xylazine (30 mg/kg), catheters (27^1/2^ gauge) were placed in the lateral tail vein followed by immediate infusion with [U-^13^C]glucose (Cambridge Isotope Laboratories, CLM-1396) at total dose of 2.48 g/kg body weight. The glucose solution was administered as a bolus (125 ul/minute over 1 minute) followed by a continuous rate of 2.5 ul/minute for 3 hours. Mice were euthanized at the end of the infusion, and plasma and liver tissues were collected and snap frozen in liquid nitrogen. Stable isotope tracing analysis was modified from a previously described protocol^45^. Metabolites were extracted from 200 mg of liver tissues using 80% ice-cold methanol after homogenization and three freeze-thaw cycles. The purified supernatants were evaporated and derivatized using TBDMS after adding 1 uL of D₂₇-myristic acid (750 ng/µL) as an internal control. One uL of the derivatized samples were injected and analyzed using an Agilent 8890 gas chromatograph coupled to an Agilent 5977C Mass Selective Detector, with data acquisition performed using MassHunter software. Data integration was conducted using El-MAVEN^46^, and the observed mass isotopologue distributions were corrected for the natural abundance of ^¹³^C using IsoCor version 2.2.2.

### Total bile acid measurement

Liver bile acids were quantified using the Total Bile Acids Colorimetric Kit (Biovision, K209-100) according to the manufacturer’s protocol. Briefly, 100 mg of liver tissue was homogenized in 500 ul of ice-cold PBS, followed by three freeze–thaw cycles to disrupt cell membranes. The homogenates were then centrifuged at 1,500 x*g* for 15 minutes at 4°C, and the supernatants were collected. 10 ul of each sample was used for the assay and bile acid levels were measured in kinetic mode at 405 nm absorbance with established standard curves.

### Magnetic resonance imaging (MRI) and analysis

Age-matched wild-type, heterozygous and homozygous *Lactb2* knockout mice were scanned at 5 weeks of age using a Bruker Biospec 94/20 MRI system (Bruker Biospin MRI GmbH, Ettlingen, Germany). Prior to scanning, mice were anesthetized in a chamber (3% Isoflurane in oxygen delivered at 0.5 L/min) and maintained using nose-cone delivery (1-2% Isoflurane in oxygen delivered at 0.2 L/min). Mice were provided with thermal support using a heated bed with warm water circulation and a physiological monitoring system to monitor breath rate. MRI was acquired with a mouse brain cryoprobe positioned over the mouse head and placed inside an 86 mm transmit/receive coil. After the localizer, T2-weighted Rapid Acquisition with Refocused Echoes (RARE) sequences were performed in the axial (TR/TE = 2000/20.4 ms, matrix size = 256 x 256, field of view = 20 mm x 20 mm, slice thickness = 0.5 mm, number of slices = 16) and coronal (TR/TE = 2500/23 ms, matrix size = 256 x 256, field of view = 20 mm x 20 mm, slice thickness = 0.5 mm, number of slices = 32) orientations. MRI image analysis was performed using 3D Slicer (Surgical Planning Laboratory) with 32 slices in the coronal direction. For measuring total brain volumes, slices from the olfactory bulb to the cerebellum were included. The brain stem was included up to and including the last slice in which the cerebellum appears. The corpus callosum was measured from every slice in which it appeared and carefully drawn for consistency between slices.

### Statistical analysis

Statistical details including sample sizes, means, and statistical significance values are indicated in the text and figure legends. Error bars in the experiments represent SD or SEM from either independent experiments or independent samples. Samples for metabolomics and GC-MS were randomized before loading to the instruments. To assess statistical significance between two groups, a two-tailed unpaired Student’s *t*-test with Welch’s correction (GraphPad 9.3) was used unless otherwise specified. Data were presented as mean +/- SD or SEM, and in all figures, the p-values were shown as: *, p<0.05; **, p<0.01; ***, p<0.001, ****, p<0.0001, n.s. not significant.

## Data availability

RNA-seq data have been deposited in the Gene Expression Omnibus database with accession number pending with NCBI. Metabolomics and lipidomics data have been provided in the Supplemental files. Further information and requests for materials and reagents should be directed to Min Ni.

---

**Extended Data Fig. 1.**
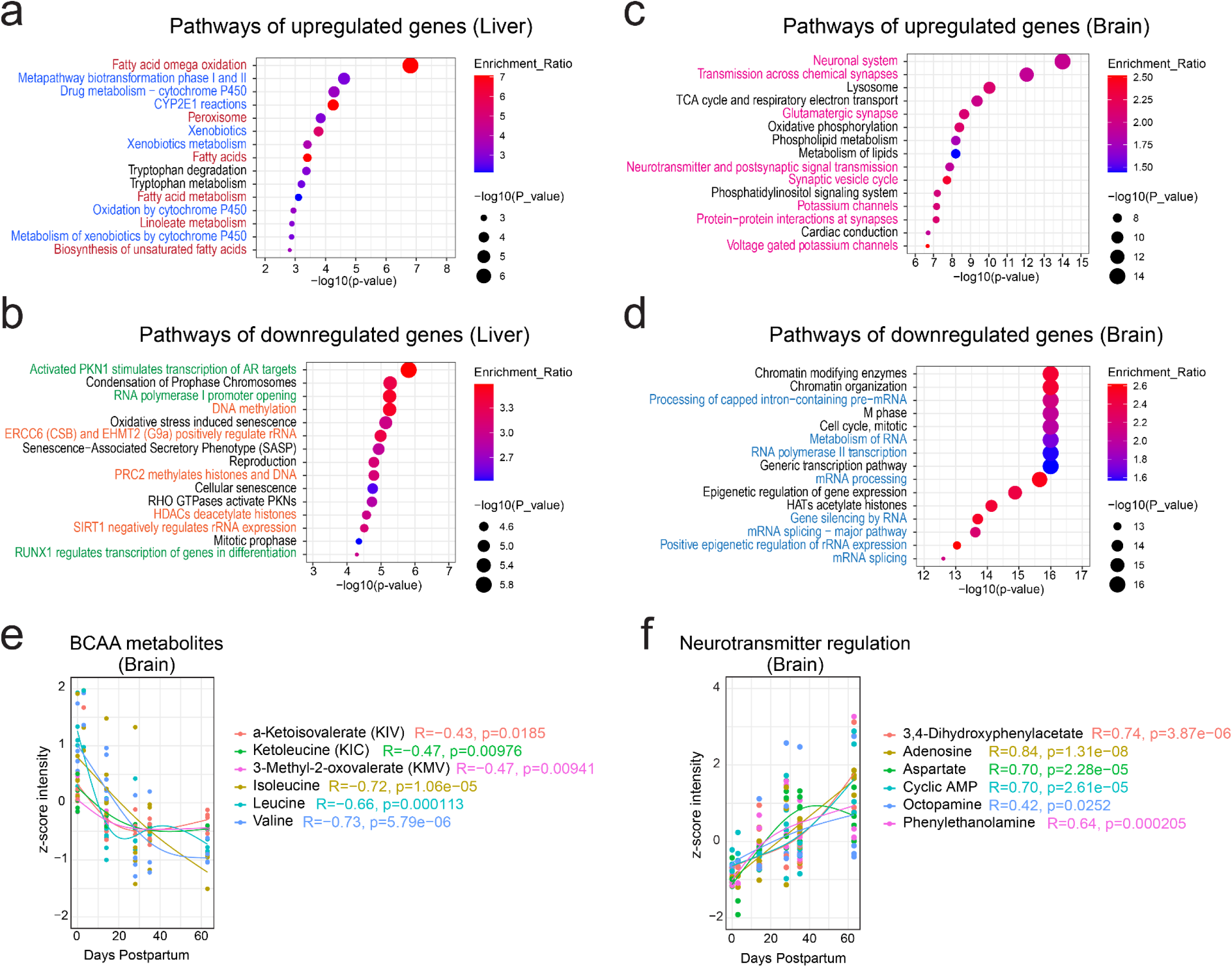
Functional analysis of organ-specific transcriptomic and metabolic changes during postnatal development. a-b. Pathway enrichment analysis of genes progressively upregulated (a) or downregulated (b) in the liver. Fatty acid metabolism (red) and detoxification pathways (blue) are highlighted in (a); transcriptional regulation (green) and epigenetic modification (orange) in (b). c-d. Pathway enrichment of genes progressively upregulated (c) or downregulated (d) in the brain. Neural function pathways are highlighted in pink (c), and RNA regulation pathways in blue (d). e. Z-score viewer of representative metabolites in branched-chain amino acid (BCAA) catabolism during postnatal brain development. f. Z-score viewer of neurotransmitters and related metabolites during postnatal brain development.

**Extended Data Fig. 2.**
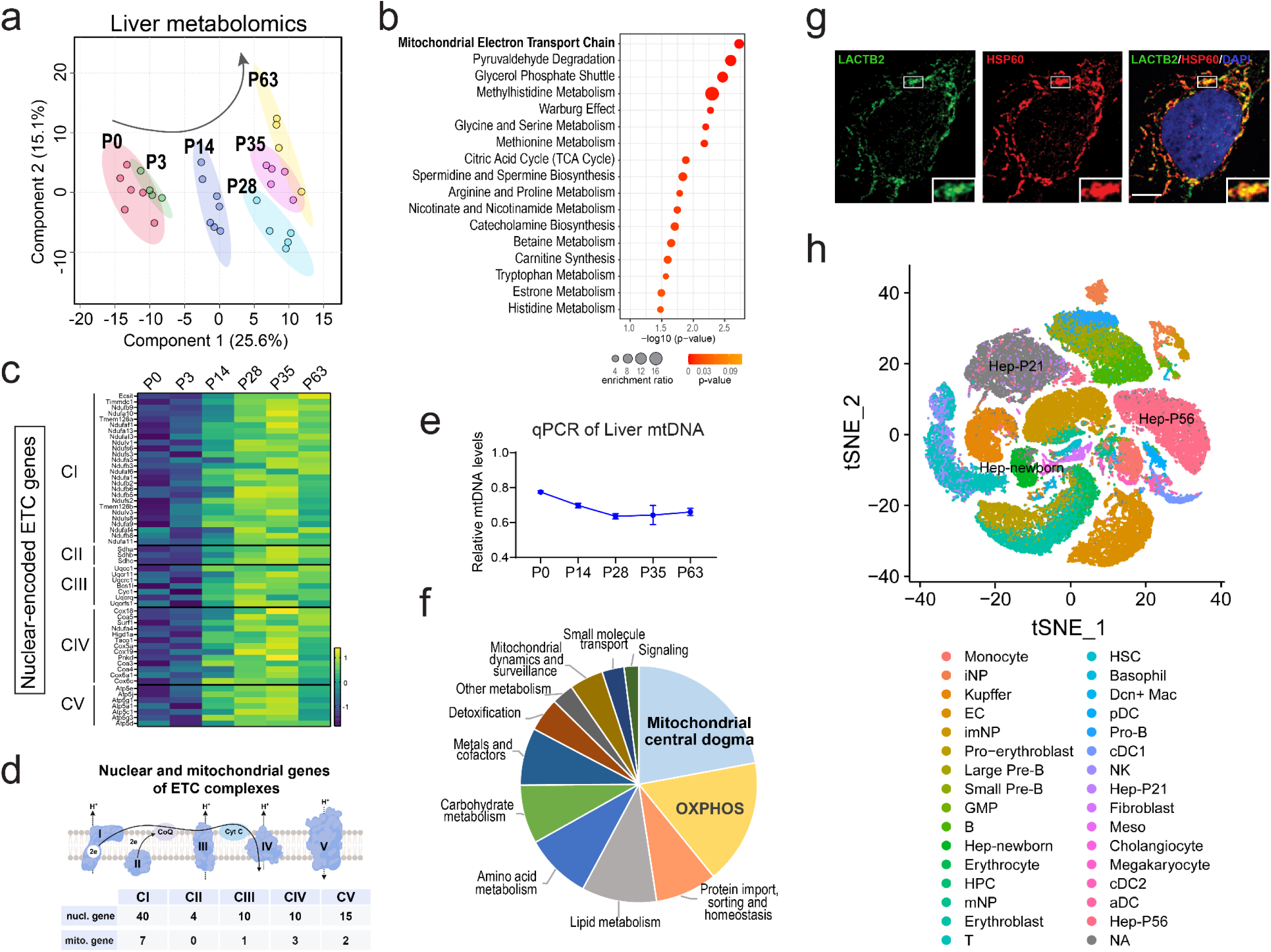
Multi-omics characterization of mitochondrial maturation in postnatal liver development. a. Partial least squares discriminant analysis (PLS-DA) of liver metabolomic profiles across six developmental stages. b. Pathway enrichment analysis of metabolites significantly altered by weaning (e.g., P3 to P35). c. Heatmap of mRNA levels for nuclear-encoded ETC genes across developmental stages. d. Schematic illustrating the contribution of nuclear and mitochondrial genomes to ETC complexes. e. Quantitative PCR analysis of mtDNA content in hepatocytes during postnatal development. f. Pie chart showing functional classification of nuclear-encoded mitochondrial genes significantly altered during liver development, based on MitoCarta 3.0 annotations. g. Immunofluorescence staining of Huh7 cells showing co-localization of LACTB2 and HSP60 in mitochondria. Scale bar, 10 um. h. tSNE plot of liver cell populations across postnatal stages from scRNA-seq data, colored by cell type clusters.

**Extended data Fig. 3.**
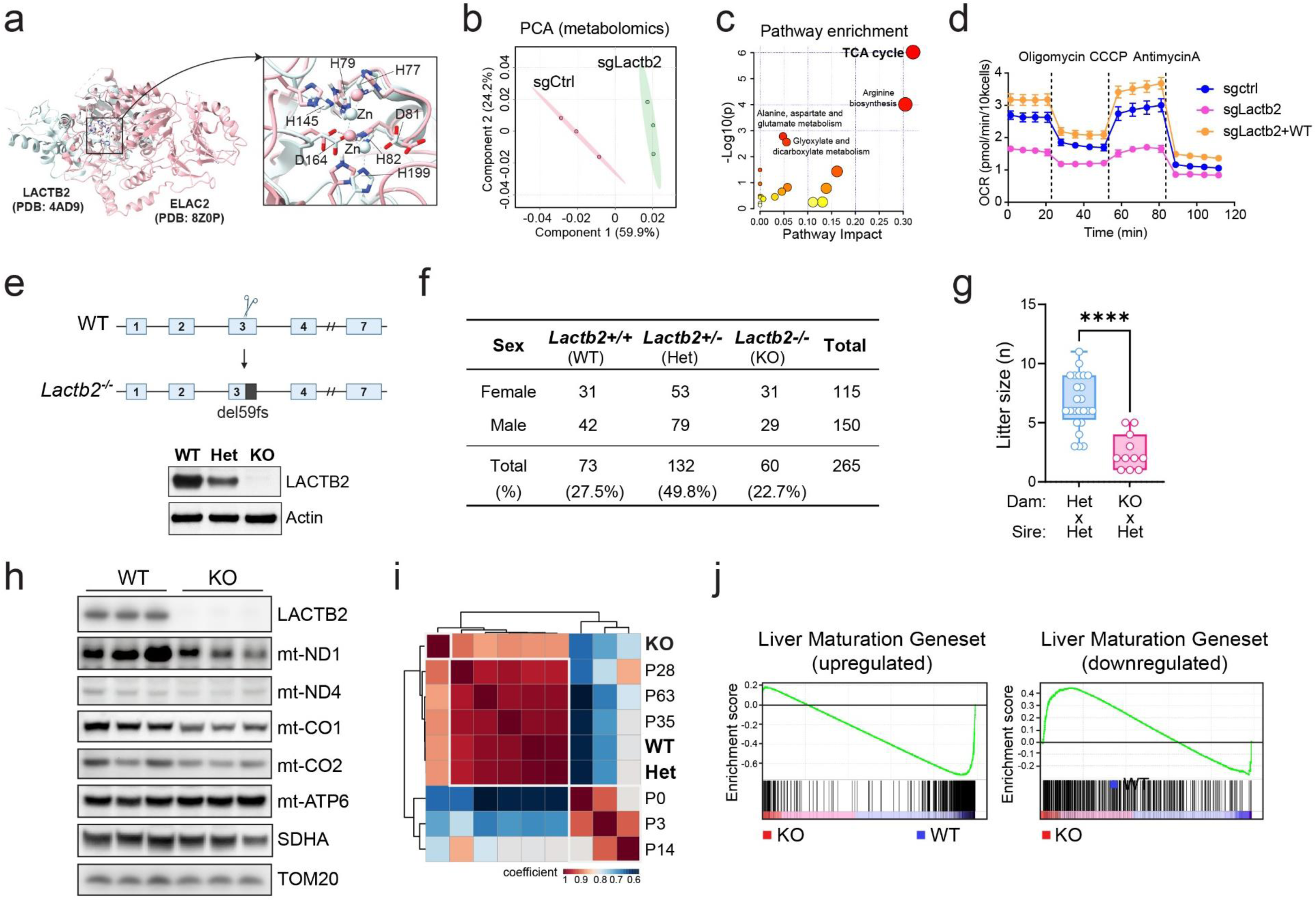
LACTB2 loss affects global liver metabolism and developmental programs. a. Superposition of human LACTB2 (PDB:4AD9) with ELAC2 (PDB: 3ZWF). A close-up of the core Zn-binding sites of two enzymes highlights key residues numbered by LACTB2’s sequence registry. b. Principal Component Analysis (PCA) of metabolomics profiles from control and LACTB2-depleted H2.35 hepatocytes. c. Metabolite set enrichment analysis (MSEA) of significantly altered metabolites in LACTB2-depleted hepatocytes versus control. d. Oxygen consumption rate (OCR) measured in control, LACTB2-depleted hepatocytes with or without re-expression of LACTB2. Results are means ± SEM from eight replicates. e. Schematic of CRISPR/Cas9-based *Lactb2* gene deletion with confirmation of protein loss in liver tissues. f. Genotype frequencies of live pups derived from *Lactb2^+/-^* intercrosses. g. Litter sizes from *Lactb2^-/-^* and *Lactb2^+/-^* dams mated with *Lactb2^+/-^* males. *P*-value by two-tailed unpaired t-test: \*\*\*\**P* < 0.0001. h. Immunoblot analysis of LACTB2 and ETC subunits in WT and KO male livers at P35 (*N* = 3 per group). i. Heatmap showing transcriptomic correlation of WT, Het and KO male livers at P35 with reference postnatal liver samples (P0-P63). Average Pearson correlation coefficients were calculated based on developmental signature genes (Supplemental Table 4). j. Gene set enrichment analysis (GSEA) of differentially expressed genes in KO versus WT male livers at P35 with gene signatures of normal postnatal liver development.

**Extended Data Fig. 4.**
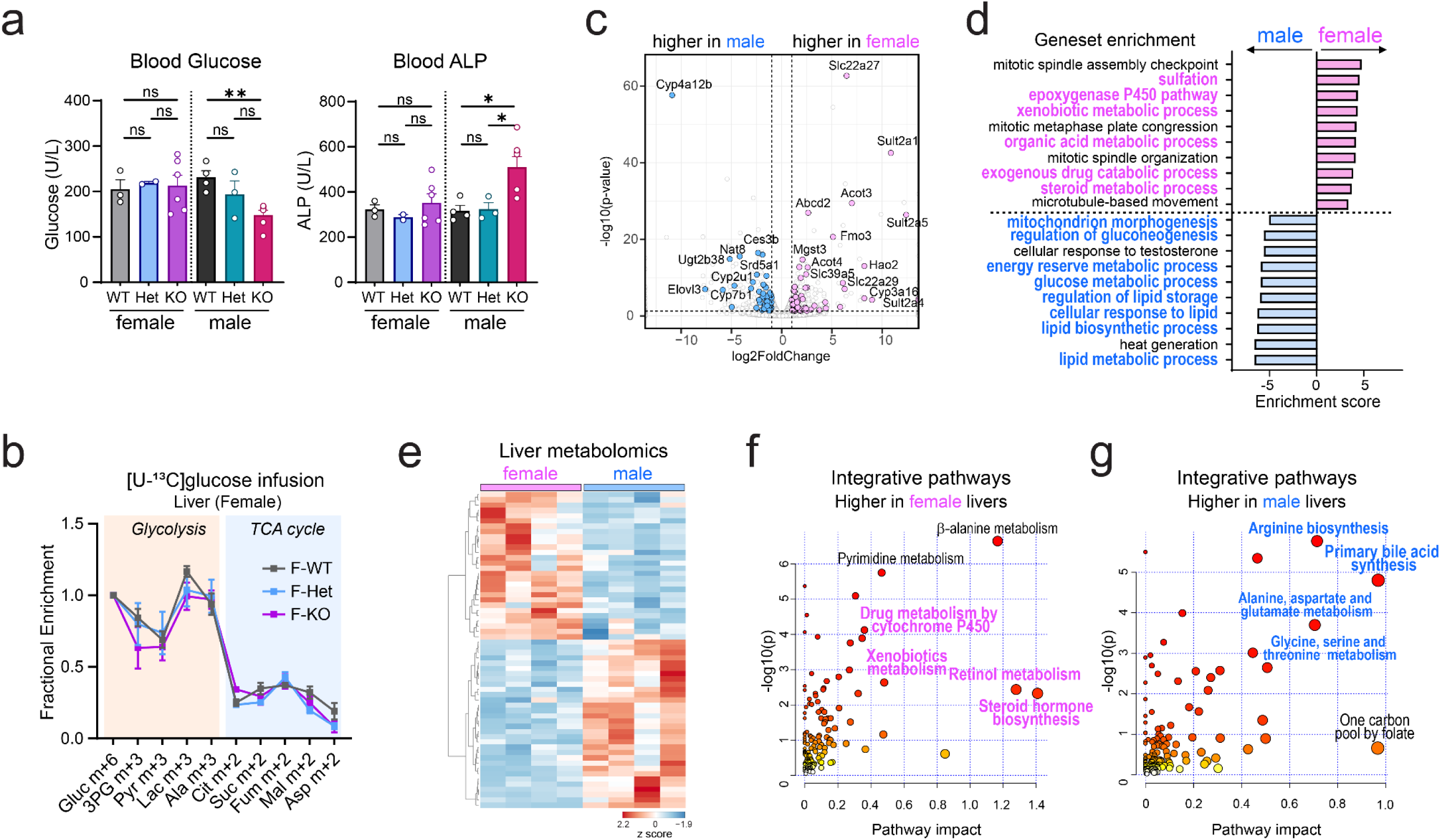
Sex-specific liver metabolism influences susceptibility to LACTB2 deficiency. a. Blood glucose and ALP levels in female and male mice with different genotypes. b. in vivo [U-^13^C]glucose tracing showing ^13^C enrichment in glycolytic and TCA cycle intermediates in female livers (*N* = 3-5 per group). c. Volcano plot of differentially expressed liver genes between female and male mice at P35, with sex-biased metabolic genes highlighted in pink for females and blue for males (*N* = 4 per group). d. Top-ranked pathways enriched for significant sex-specific genes in the liver. Metabolic pathways upregulated in females are highlighted in pink, and those upregulated in males are highlighted in blue. e. Heatmap of liver metabolites with variable importance in the projection (VIP) scores ≥ 1.0 comparing females and males at P35. f-g, Integrative pathway enrichment of differential liver metabolites and metabolic genes between female and male mice at P35.

**Extended Data Fig. 5.**
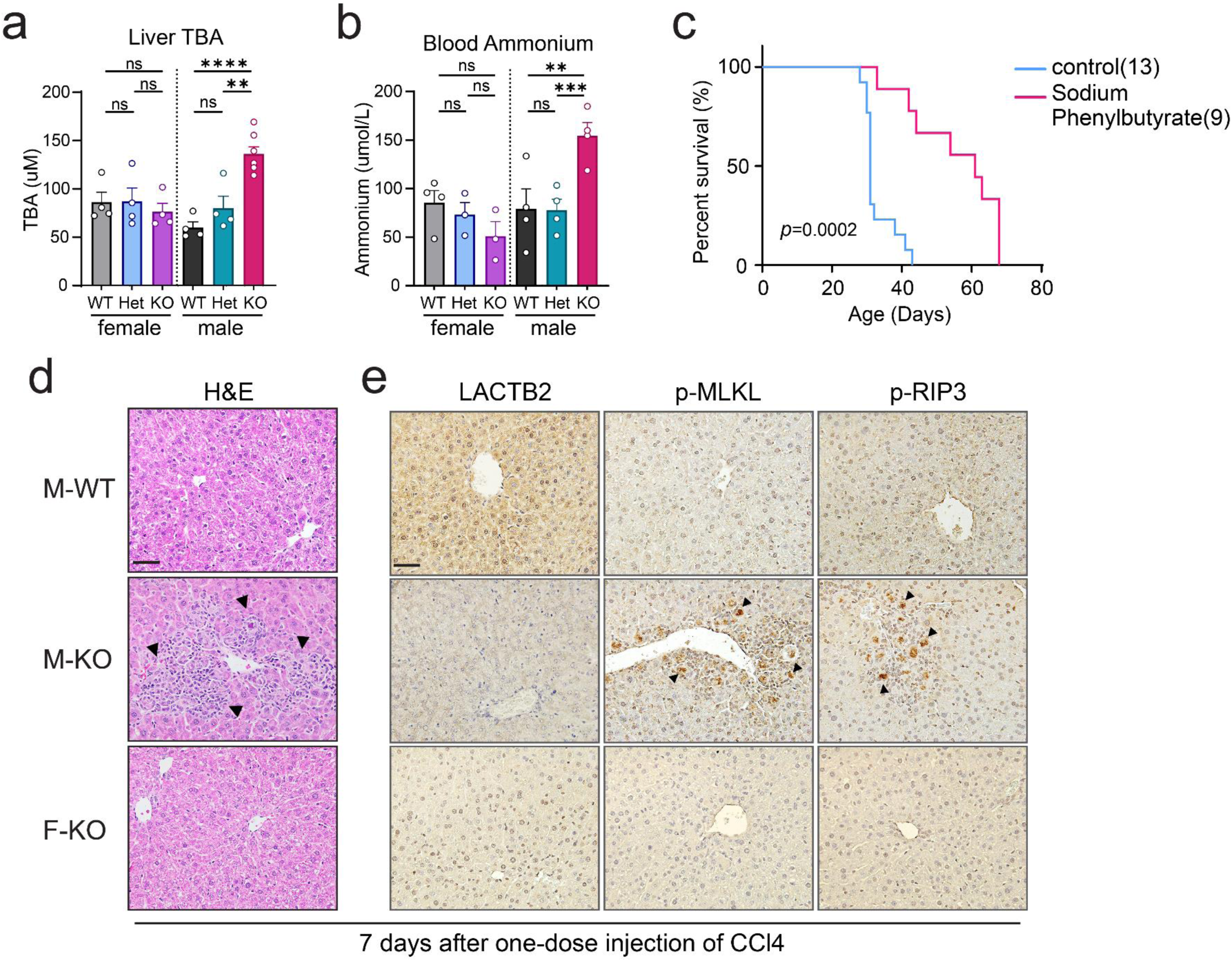
LACTB2-deficient males exhibit impaired hepatic detoxification capacity. a. Total bile acid (TBA) levels in liver tissue from the indicated female and male groups (*N* = 4-7 mice per group). Data analyzed by two-tailed unpaired t-test. \**P* < 0.05, \*\**P* < 0.01, \*\*\**P* < 0.001, n.s. not significant. b. Blood ammonium levels in female and male groups of all three genotypes at P35 (*N* = 3-5 mice per group). *P* values by two-tailed unpaired t-test: \*\**P* < 0.01, \*\*\**P* < 0.001, \*\*\*\**P* < 0.0001, ns, not significant. c. Survival of KO males with or without sodium phenylbutyrate treatment. *P* value by log-rank Mantel-Cox test. d. H&E staining of FFPE liver samples from WT and KO mice 7 days after a single hepatotoxic dose of carbon tetrachloride (CCl4; 5 ml/kg body weight). *N* = 3 mice per group. Scale bar, 50 um. e. IHC staining of LACTB2 and necroptosis markers p-MLKL and p-RIP3 in livers from WT and KO mice 7 days post-CCL4 injection. *N* = 3 mice per group. Scale bar, 50 um.

**Extended Data Fig. 6.**
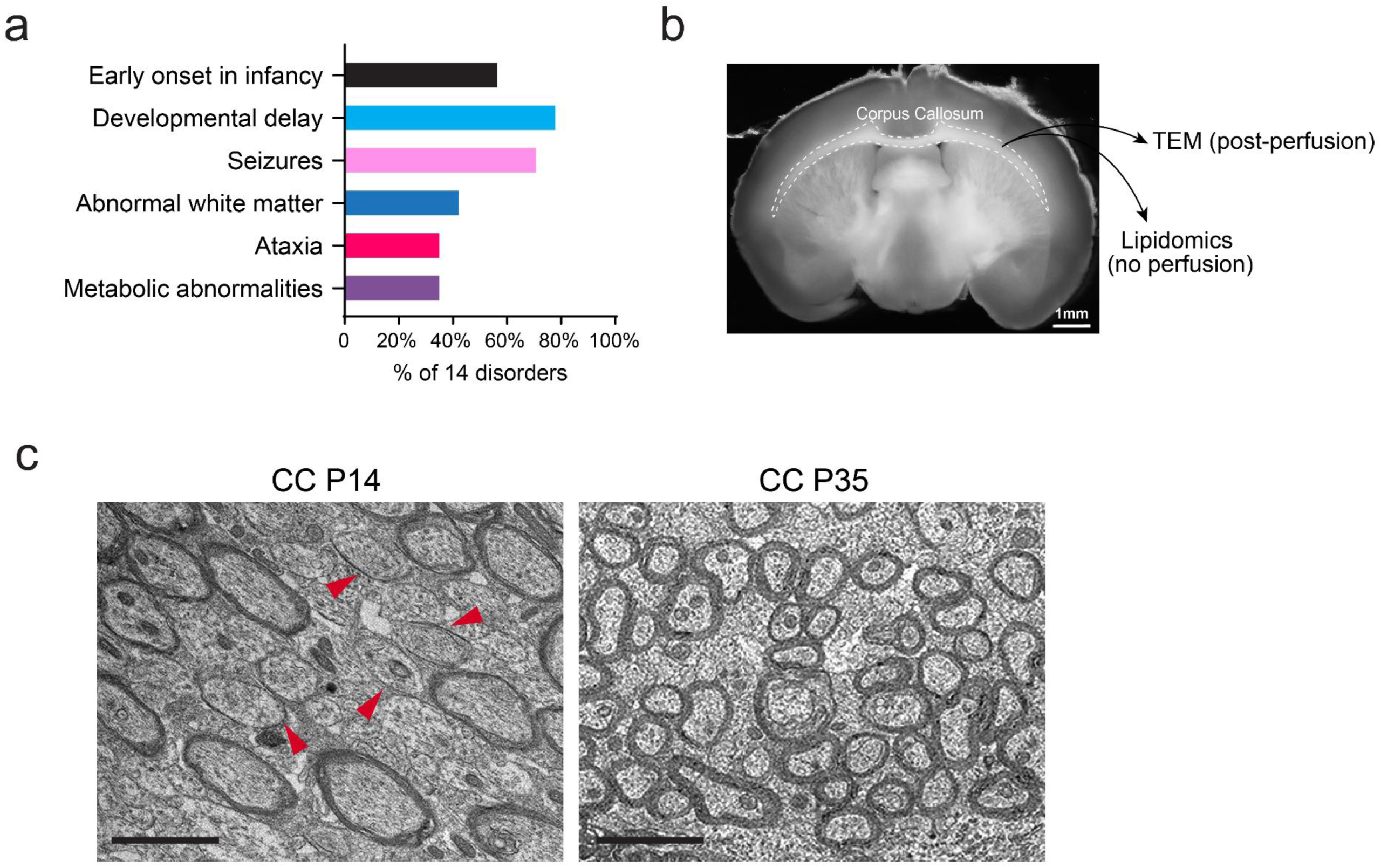
Neurological involvement associated with LACTB2 deficiency. a. Neurological and metabolic phenotypes linked to disease-associated genes that are significantly downregulated in KO male brains compared to WT. Clinical manifestations are annotated based on OMIM clinical synopses. b. Illustration of the mouse corpus callosum in a coronal brain section. For TEM, tissues were collected after perfusion fixation. For lipidomics, fresh non-perfused corpus callosum tissue was dissected and processed. c. TEM images showing corpus callosum myelin in male brains at P14 and P35. Red arrowheads indicate immature or newly formed myelin sheaths. Scale bars, 2 µm.

